# Salivary gland macrophages assist tissue-resident CD8^+^ T cell immune surveillance

**DOI:** 10.1101/723791

**Authors:** B. Stolp, F. Thelen, X. Ficht, L. M. Altenburger, N. Ruef, V. V. G. K. Inavalli, P. Germann, N. Page, F. Moalli, A. Raimondi, K. A. Keyser, S. M. Seyed Jafari, F. Barone, M. S. Dettmer, D. Merkler, M. Iannacone, J. Sharpe, C. Schlapbach, O. T. Fackler, U. V. Nägerl, J. V. Stein

## Abstract

Tissue macrophages and tissue resident memory CD8^+^ T cells (T_RM_) play important roles for pathogen sensing and rapid protection of barrier tissues. To date, it is incompletely understood how these two cell types cooperate for efficient organ surveillance during homeostasis. Here, we used intravital imaging to show that T_RM_ dynamically crawled along tissue macrophages in murine submandibular salivary glands (SMG) during the memory phase following a viral infection. *Ex vivo* confined SMG T_RM_ integrated an unexpectedly wide range of migration modes: in addition to chemokine-and adhesion receptor-driven motility, SMG T_RM_ displayed a remarkable capacity of autonomous motility in the absence of chemoattractants and adhesive ligands. This unique intrinsic SMG T_RM_ motility was transmitted by friction and adaptation to microenvironmental topography through protrusion insertion into permissive gaps. Analysis of extracellular space in SMG using super-resolution shadow imaging showed discontinuous attachment of tissue macrophages to neighboring epithelial cells, offering paths of least resistance for patrolling T_RM_. Upon tissue macrophage depletion, intraepithelial SMG T_RM_ showed decreased motility and reduced epithelial crossing events, and failed to cluster in response to local inflammatory chemokine stimuli. In sum, our data uncover a continuum of SMG T_RM_ migration modes and identify a new accessory function of tissue macrophages to facilitate T_RM_ patrolling of the complex exocrine gland architecture.

**One sentence summary:** Combined *in vitro* and *in vivo* imaging of salivary gland-resident tissue memory CD8^+^ T cells (T_RM_) uncovers their unique migratory behavior and describes a novel accessory function of tissue macrophages to assist T_RM_ surveillance.

## Introduction

Naïve CD8^+^ T cells (T_N_) continuously traffic through lymphoid tissue such as peripheral lymph nodes (PLN) and spleen, where they screen antigen presenting dendritic cells (DCs) for the presence of cognate peptide-MHC (pMHC) complexes. Intravital two-photon microscopy (2PM) of peripheral lymph nodes (PLN) uncovered a high amoeboid-like T_N_ motility of 12-15 µm/min (*1–4*), facilitating their search for rare cognate pMHC-presenting DCs interspersed on a 3D stromal scaffold of fibroblastic reticular cells (FRC) (*5–10*). Intranodal motility is mediated by the CCR7 ligands CCL19 and CCL21 that drive F-actin polymerization at the leading edge in a Gαi-dependent manner to generate a retrograde cortical actin flow. Cortical actin flow is conveyed by the integrin LFA-1 into forward movement without generating substantial substrate adhesion (*11–15*).

During viral infections, effector CD8^+^ T cells (T_EFF_) generated in reactive lymphoid tissue disseminate into non-lymphoid tissues (NLT) including gut, lung, genitourinary tract, and skin to eliminate infected cells. After clearance of viral antigens, part of T_EFF_ differentiate into central memory T cells (T_CM_) and continue to patrol lymphoid organs, while others stably reside in NLT and to a minor extent in lymphoid tissue as non-recirculating, self-renewing tissue-resident memory T cells (T_RM_). T_RM_ in gut, skin, and genitourinary tract act as “first line” sentinels that eliminate infected cells and trigger an organ-wide alert status through cytokine secretion upon pathogen re-encounter (*16–21*). The scanning behavior of epidermal T_RM_ has been extensively studied. These cells display characteristically elongated, dendritic shapes and move in a Gαi-dependent manner with speeds of 1-2 µm/min in proximity to the extracellular matrix (ECM)-rich basement membrane (BM) separating epidermis from dermis, i.e. in plane with the bottom keratinocyte layer (*22, 23*). Upon pathogen reencounter, epidermal CD8^+^ T cells follow local chemokine signals to accumulate around infected cells (*24*). CD8^+^ T cell accumulation is considered critical for cooperative elimination of infected stromal cells through repeated cytotoxic attacks (*25*). Similarly, CD8^+^ T_RM_ of the small intestine continuously patrol the absorptive epithelial layer (*26*).

T_RM_ are also present in exocrine glands of the head and neck region, including submandibular salivary glands (SMG). Salivary glands are targeted by several bacteria and viruses including human beta- and gamma-herpesviruses, which can cause disease, mostly in immunocompromised individuals (*27, 28*). Similar to skin and gut, SMG contains epithelial tissue basally anchored onto connective tissue. However, while skin and gut 3D geometry is evenly layered, the SMG epithelium has an arborized structure, with acini secreting saliva into intermediate and collecting ducts. The glandular epithelium is separated by a BM from the supporting interstitium containing blood and lymphatic vasculature, fibroblasts and tissue macrophages (*29*). In tissue sections, most CD8^+^ T_RM_ in SMG are localized within the abundant acini and ducts, implying a mechanism that allows T_EFF_ arriving in interstitial venules to cross the BM below the epithelial compartment and develop into memory T cells (*30, 31*). During acute inflammation of NLT, CD8^+^ T cell recruitment is driven by chemokines and adhesion receptors (*32*). In contrast, the cellular dynamics of homeostatic T_RM_ surveillance in SMG after viral infection and the involvement of tissue macrophages in this process have not been explored to date.

Here, we used intravital microscopy of mouse SMG in the memory phase following a systemic viral infection to uncover a high baseline motility of T_RM_, which often followed tissue macrophage topology. *Ex vivo*, confinement alone in the absence of chemoattractants and adhesion receptors was sufficient to induce SMG T_RM_ migration through friction- and protrusion-insertion-driven motility, which was further tuned by chemokines and adhesion molecules. Using super-resolution microscopy to explore extracellular space distribution in SMG, we observed discontinuous attachment of tissue macrophages to surrounding epithelium, offering paths of least resistance to migrating T_RM_. Accordingly, tissue macrophage depletion resulted in a significant disruption of T_RM_ patrolling behavior. Taken together, our data uncover a new accessory role for tissue macrophages to enable T_RM_ surveillance of salivary glands. Our observations suggest a continuum of chemokine- and adhesion receptor-dependent and -independent migration modes and topographic features facilitating this task.

## Results

### Systemic viral infection leads to the establishment of T_RM_ in salivary glands

We used a viral infection model for a comparative analysis of CD8^+^ T cell populations in lymphoid tissue and SMG (**Fig. 1A and Fig. S1A**). Systemic infection with lymphocytic choriomeningitis virus (LCMV)-OVA, a replication-competent, attenuated LCMV mutant expressing ovalbumin (OVA) as model antigen (*33*), led to transient and low viral titers in spleen on day 3 p.i. that remained below the detection limit in PLN and SMG (**Fig. S1B**). Adoptively transferred GFP^+^ OT-I CD8^+^ TCR tg T cells (which recognize the OVA_257-264_ peptide in the context of H2-K^b^) (*34*) underwent a prototypic expansion – contraction kinetic in spleen and PLN over the course of 30 days (**Fig. S1, C and D**). Despite the lack of detectable viral titers, OT-I T cells accumulated in SMG from day 6 p.i. onwards, with a stable population maintained until at least day 30 p.i. (**Fig. S1, C and D**). By day 30 p.i., OT-I T cells isolated from SMG but not PLN or spleen showed increased expression of CD103 and CD69, while losing the KLRG-1^+^ population present on day 6 p.i. (**Fig. S1, E and F**). SMG OT-I T cells also upregulated PD-1 and CD44 surface levels (not shown), supporting the observation that most SMG CD8^+^ T cells had developed into *bona fide* T_RM_ at day 30 p.i. (*35*). Memory OT-I CD8^+^ T cells isolated from PLN were approximately 65% CD62L^+^ CD44^high^ T_CM_, with the remaining population being CD62L^-^ CD44^high^ memory T cells. To take this heterogeneity into account, we refer to memory T cells isolated from PLN as T_PLN-M_.

**Fig. 1.**
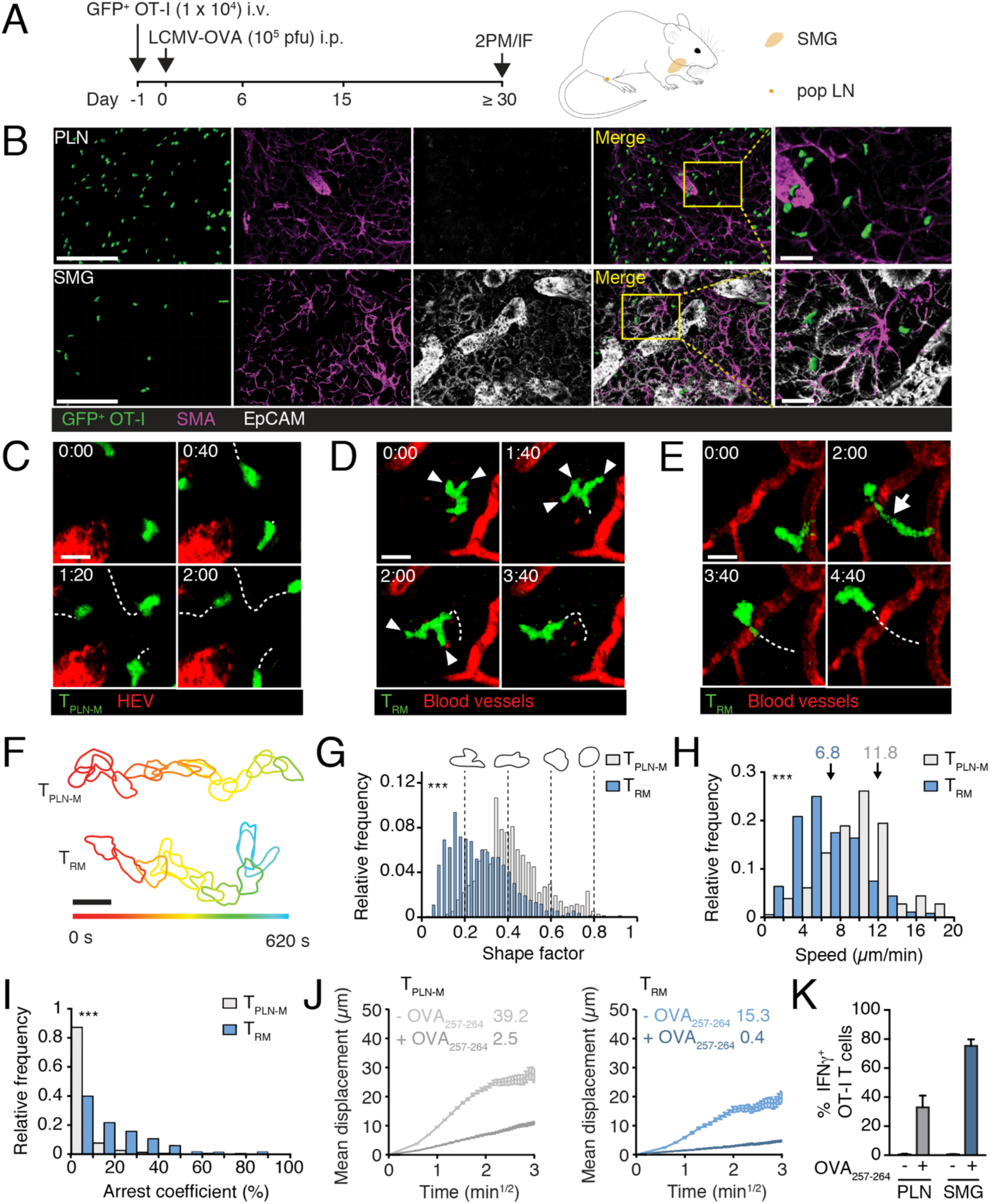
Dynamic motility parameters of memory CD8^+^ T cells in PLN versus SMG. **A.** Experimental layout for CD8^+^ T cell analysis in SMG and PLN. **B.** Immunofluorescent sections of GFP^+^ OT-I T cells in PLN and SMG in memory phase (≥ day 30 p.i.). Scale bar, 100 µm (left panels) and 20 µm (right panel). **C.** Time-lapse 2PM image sequences showing OT-I CD8^+^ T_PLN-M_ cell motility in PLN in memory phase (≥ day 30 p.i.). **D and E.** Time-lapse 2PM image sequences showing OT-I CD8^+^ T_RM_ cell motility in SMG in memory phase (≥ day 30 p.i.). Arrowheads indicate protrusions (**D**) and the arrow indicates squeezing behavior (**E**) of OT-I CD8^+^ T_RM_. Scale bar in C-E, 10 µm. Time in min:s. **F**. Time-coded shapes of exemplary T_PLN-M_ and T_RM_ tracks. **G.** Shape factor distribution of T_PLN-M_ and T_RM_ with exemplary cell shapes. **H.** Speed frequency distribution of OT-I CD8^+^ T cells in PLN and SMG. Arrows indicate median values (µm/min). **I.** Arrest coefficient frequency distribution of OT-I CD8^+^ T cells in PLN and SMG (cut-off < 2.5 µm/min). **J.** Mean displacement versus time of OT-I T_PLN-M_ (left) and T_RM_ (right) before and after OVA_257-264_ injection with motility coefficients (μm^2^/min). **K.** IFN-γ expression in OT-I T_PLN-M_ and T_RM_ 24 h after OVA_257-264_ injection (mean ± SD). Data in G are from 2-3 independent experiments and 3 mice total for each group. Data in H and I are pooled from 5 to 6 mice from 4 independent experiments with at least 194 tracks analyzed per organ. Data in J are pooled of 3-4 mice from 2 independent experiments. Data in K show one of two independent experiments. Data in G and I were analyzed with Mann-Whitney-test and data in H with Student’s t-test. ***, p < 0.001.

T_RM_ establishment in SMG was also observed at ≥ 30 day after systemic VSV-OVA infection (not shown). In addition, we detected memory P14 CD8^+^ TCR tg T cells (which recognize the LCMV epitope gp_33-41_ in the context of H2-D^b^) (*36*) in SMG after infection with the LCMV Armstrong strain (**Fig. S2, A and B**). In sum, systemic viral infection led to the recruitment and retention of CD8^+^ T cell populations in SMG, even in the absence of detectable viral titers in this organ.

### SMG T_RM_ migration is characterized by dynamic cell shape changes

We next determined the localization of T_PLN-M_ and T_RM_ in their target organs during the memory phase. Tissue sections showed that most GFP^+^ OT-I T cells in PLN and SMG were dispersed evenly in the tissue at day 30 p.i. (**Fig. 1B**). In PLN, OT-I T_PLN-M_ cells localized mainly with smooth muscle actin (SMA)^+^ FRCs, whereas most OT-I T_RM_ cells in SMG were within or adjacent to EpCAM^+^ acini and ducts (**Fig. 1B**). We developed a sequential surgery method to visualize homeostatic tissue surveillance of T_PLN-M_ and T_RM_ in PLN and SMG, respectively, in the same host by 2PM (*37*). T_PLN-M_ displayed characteristic amoeboid shapes and moved with high speeds comparable to T_N_ (11.8 ± 4.0 µm/min, median ± SD) (**Fig. 1, C and F to H; movie S1**). Compared to T_PLN-M_, SMG T_RM_ displayed more pronounced shape changes, with several protrusions probing the microenvironment during migration, at times with thin and elongated cell bodies (**Fig. 1, D to G; movie S2**). While T_PLN-M_ and T_RM_ covered large distances throughout the observation period of intravital imaging sequences (20-60 min), both populations differed in their speed and arrest coefficients, i.e. percentage of track segments with speeds < 2.5 µm/min. Thus, SMG T_RM_ were significantly slower than T_PLN-M_ (**Fig. 1H**) and had higher arrest coefficients (**Fig. 1I**). Nonetheless, SMG T_RM_ retained a relatively high motility coefficient, which is a proxy of a cell’s ability to scan the environment during random migration, of > 15 µm^2^/min (**Fig. 1J**). Accordingly, their median speed of 6.8 ± 3.4 µm/min was notably higher than values reported for epidermal T_RM_ (1-2 µm/min) (*22*), with some cells achieving speeds of > 12 µm/min. Both T_PLN-M_ and SMG T_RM_ retained a fast response to antigenic stimulation, as systemic administration of cognate peptide resulted in immediate arrest and secretion of IFN-γ (**Fig. 1, J and K; Fig. S2, C and D**).

We measured similar speeds for GFP^+^ P14 T_RM_ in SMG before and after cognate peptide administration (**Fig. S2**, A to D). Furthermore, GFP^+^ OT-I T cells patrolled the structurally comparable lacrimal gland (LG) in the same speed range (7.6 ± 4.3 µm/min; median ± SD; n = 255 tracks), suggesting that migration parameters of T_RM_ patrolling exocrine glands during homeostasis are independent of TCR specificity and reflect tissue properties. In sum, our data uncover a remarkably fast motility of exocrine gland-resident CD8^+^ T cells, which was characterized by dynamic shape changes.

### T_RM_ migrate along tissue macrophages during SMG surveillance

To explore the microenvironmental context of exocrine gland T_RM_ migration, we used a CD11c-YFP reporter strain that labels SMG CD64^+^ F4/80^+^ tissue macrophages (*29*). CD11c-YFP^+^ cells were also positive for the macrophage marker Iba-1, whereas some Iba-1^+^ cells were CD11c-YFP^low/negative^, indicating that most but not all tissue macrophages were labeled in CD11c-YFP mice (**Fig. S3, A and B**). Confocal analysis of thick tissue sections showed that CD11c-YFP^+^ tissue macrophages extended numerous protrusions from their cell bodies throughout the SMG tissue and were located within EpCAM^+^ ducts and acini, as well as SMA^+^ perivascular structures of the interstitium (**Fig. S3C**). Most macrophage protrusions were phosphotyrosine-positive (**Fig. S3, D and E**) and enriched in F-actin (not shown), suggesting the presence of podosomes or focal adhesions at these sites.

To assess the spatial relationship between tissue macrophages and T_RM_, we transferred GFP^+^ OT-I T cells into CD11c-YFP recipients one day prior to infection with LCMV-OVA and analyzed tissue sections by confocal microscopy in memory phase (≥ day 30 p.i.). We observed a striking spatial proximity of T_RM_ and tissue macrophages in SMG, with approximately 70% of OT-I T cells directly in contact with CD11c-YFP^+^ cells (**Fig. 2, A and B; movie S3**). The close spatial association between tissue macrophages and T_RM_ was confirmed by correlative light and electron microscopy imaging, with both cell membranes adjacent to each other (**Fig. 2C**). Electron microscopy images also highlighted the compact tissue structure of SMG, with tight junctions of acinar and ductal epithelium surrounded by a dense ECM (**Fig. 2D**).

**Fig. 2.**
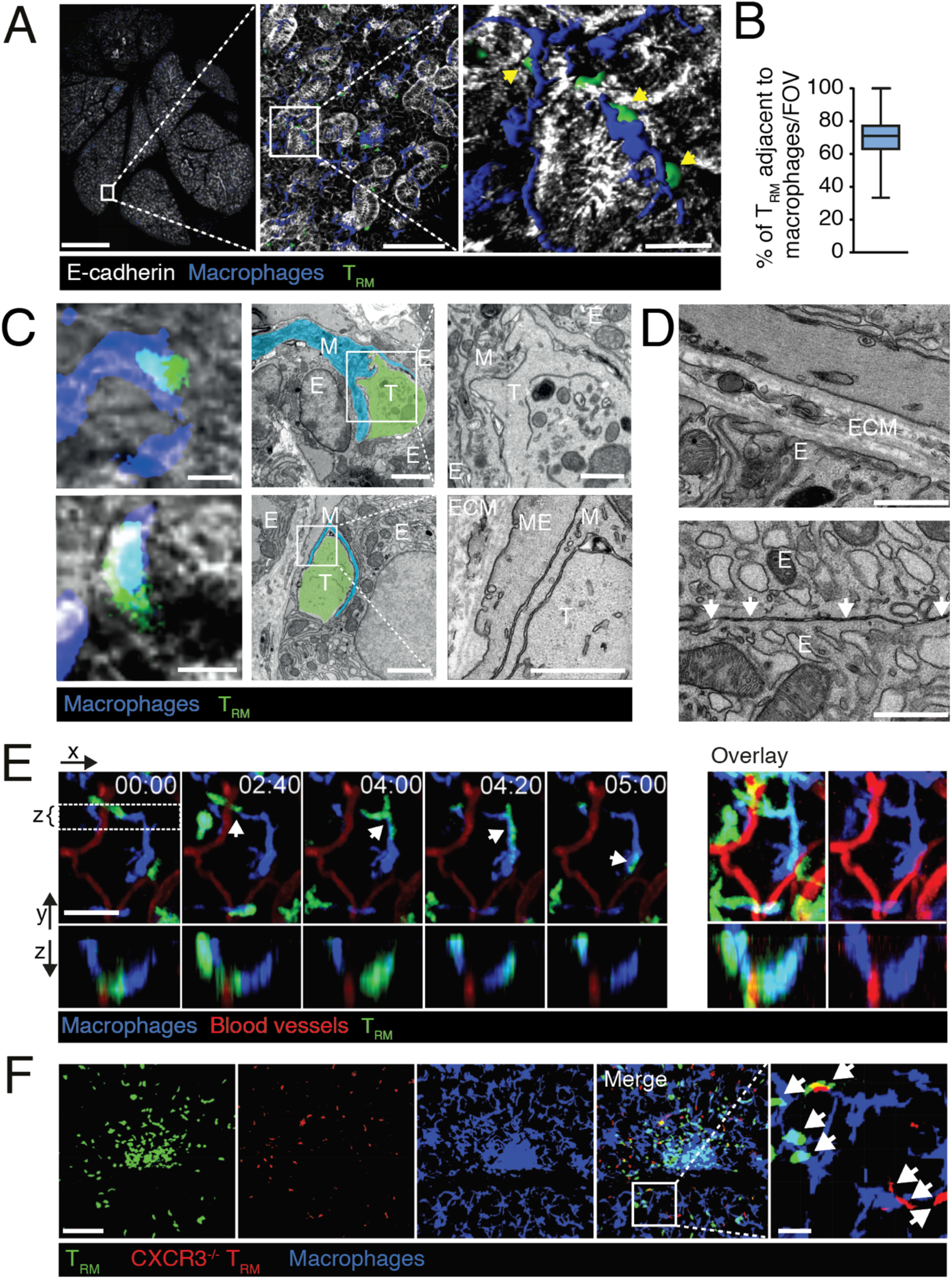
SMG T_RM_ move alongside tissue macrophages. **A.** Immunofluorescent section showing localization of SMG T_RM_ adjacent to tissue macrophages (arrows). Scale bars, 1 mm (left), 100 µm (middle) and 20 µm (right). **B.** Percent of SMG T_RM_ adjacent to tissue macrophages. Data are pooled from 105 FOV with a total of 3270 T_RM_ and shown as box and whisker graph with 2.5 – 97.5 percentiles. **C.** Correlative light and electron microscopy sections (left; confocal image; middle and right, TEM image) showing close spatial association of SMG T_RM_ and tissue macrophages. M, tissue macrophages; E, epithelial cell; ME, myoepithelial cell; ECM, extracellular matrix. Scale bar, 5 (left), 2 (middle) and 1 µm (right). **D.** TEM images showing attachment of epithelial cells to ECM (top) and through intercellular junctions (white arrows; bottom). Scale bar, 800 nm. **E.** 2PM time-lapse image sequence showing overlap of OT-I T_RM_ tracks with tissue macrophages in SMG in memory phase (≥ day 30 p.i.). Scale bar, 20 µm. Time in min:s. The right panels show the time accumulated overlays of images with or without OT-I T_RM_. **F.** Immunofluorescent section of WT and CXCR3^-/-^ OT-I T cells and macrophages. Magnified image shows association of CXCR3^-/-^ OT-I T_RM_ to tissue macrophages (arrows). Scale bar, 100 µm (left) and 20 µm (right).

Their spatial proximity to SMG macrophages in tissue sections raised the question whether patrolling T_RM_ migrate alongside macrophages. 2PM imaging of GFP^+^ T_RM_ in LCMV-OVA memory phase CD11c-YFP recipients indeed confirmed that T_RM_ crawled along CD11c-YFP^+^ macrophages during most of the observation period, with T_RM_ shapes often closely matching the underlying macrophage topology. This was particularly evident along thin macrophage protrusions, which T_RM_ often followed (**Fig. 2E; movie S4**). At the same time, T_RM_ protrusions occasionally detached from macrophages, apparently scanning the surrounding environment. Accordingly, we identified occasional T_RM_ track segments which were not associated with tissue macrophages. T_RM_ speeds were slightly elevated when in contact with tissue macrophages than when not (7.0 ± 5.3 versus 6.1 ± 4.8 µm/min; p < 0.001). Occasionally, we observed small T_RM_ clusters around tissue macrophages. Adoptively transferred CXCR3^-/-^ T_RM_ failed to accumulate at tissue macrophage clusters, suggesting the existence of local CXCL9/CXCL10 “hotspots” at these sites (**Fig. 2F**).

We examined whether the noticeable proximity between tissue macrophages and T_RM_ also occurred in other exocrine glands and species. A comparable association of T_RM_ and tissue macrophages was observed in LG after LCMV-OVA infection (**Fig. S3F**). Furthermore, CD3^+^ T cells colocalized with CD68^+^ macrophages in human parotid gland sections, both as dispersed individual cells and in clusters (**Fig. S3, G and H**). Taken together, SMG T_RM_ colocalized with and moved alongside tissue macrophages during homeostatic tissue patrolling of several exocrine glands.

### SMG T_RM_ motility is induced by confinement and can be tuned by external factors

The close proximity of T_RM_ to tissue macrophages *in vivo* prompted us to examine the molecular factors involved in this interaction. We performed quantitative PCR analysis of cytokine and chemokine expression by CD11c-YFP^+^ cells sorted from SMG in steady-state, acute (day 6 p.i.) and memory phase (day 30 p.i.) of LCMV-OVA infection. We observed detectable mRNA levels of the cytokines IL-1 and TNF, as well as the chemokines CCL3, CCL4, CXCL2, CXCL9, CXCL10 and CXCL16 (not shown). Expression levels were similar at all time points analyzed, reflecting the lack of detectable viral spread to this organ (**Fig. S1B**). Given the expression of promigratory chemokines and adhesion receptors including ICAM-1 on tissue macrophages (*38*), we examined their influence on T_RM_ migration parameters. To this end, we employed under agarose assays that allow to precisely control environmental factors and provide the confinement required for T cell motility (**Fig. 3A**) (*15*). To benchmark our system, we transferred T_N_ on CCL21 - and ICAM-1-coated plates as surrogate lymphoid tissue microenvironment. We observed high chemokinetic T_N_ motility with similar speeds as measured *in vivo* (13.3 ± 5.9 µm/min) (**Fig. 3, B and C; movie S5**) (*14, 15*). T_RM_ showed a high motility (11.4 ± 3.0 µm/min) when migrating on CXCL10 + CXCL12- and ICAM-1-coated plates, which was only slightly lower than that of T_N_ (**Fig. 3C; movie S6**). These observations show that SMG T_RM_ respond to presence of chemokines and adhesion molecules with high speeds.

**Fig. 3.**
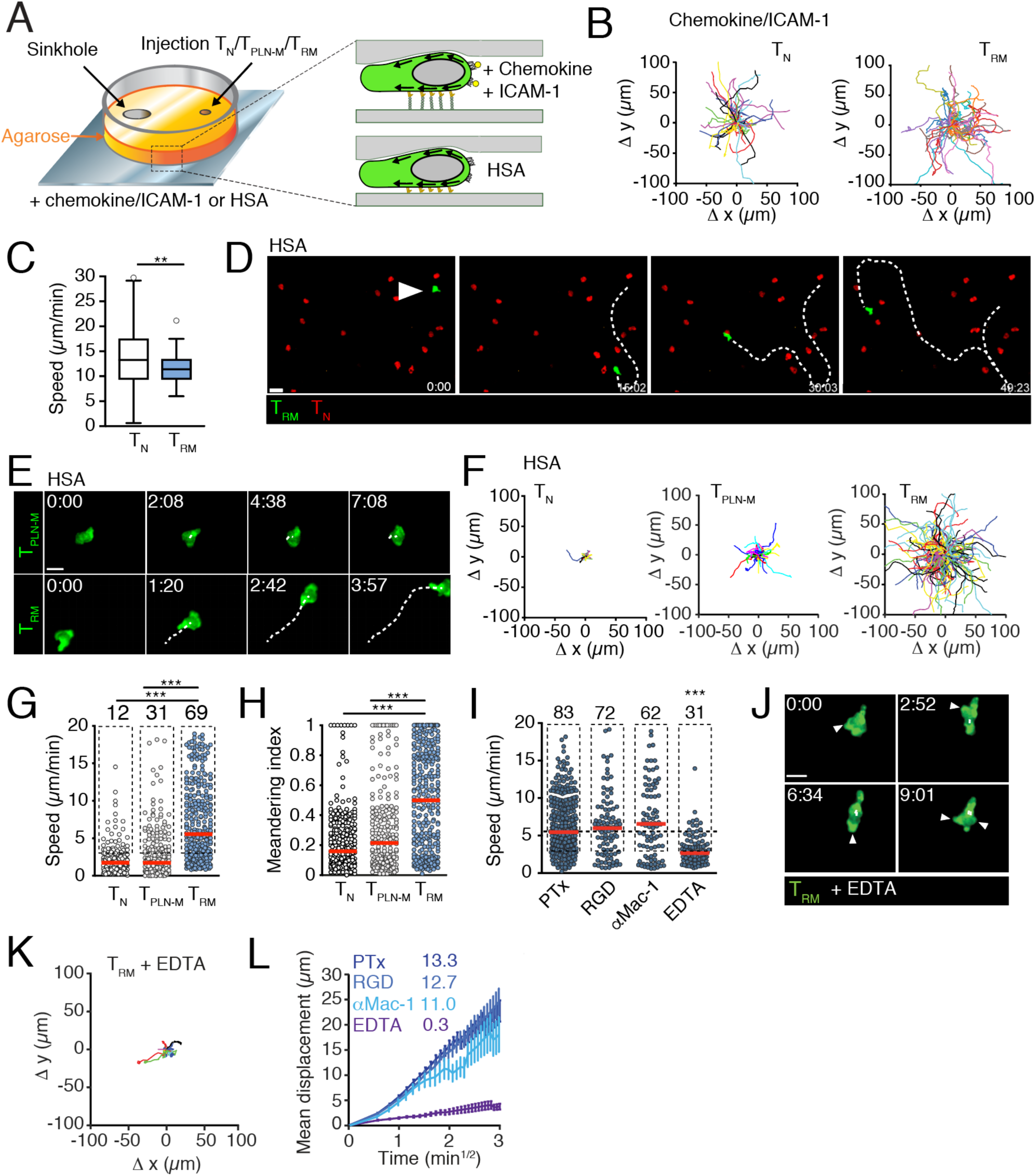
Confinement induces SMG T_RM_ motility through chemokine- and adhesion-mediated signals and bivalent cation-dependent friction. **A.** Experimental layout of under agarose assay. Arrows indicate F-actin flow. **B.** Representative T_N_ (n = 75) and T_RM_ (n = 58) tracks in presence of chemokine and ICAM-1. **C.** Speeds of T_N_ and T_RM_. Data are presented as Tukey box and whiskers plot. **D.** Time-lapse image sequence showing T_RM_ motility among immotile T_N_. T_RM_ displacement shown by segmented line. Scale bar, 20 μm. Time in min:s. **E.** Time-lapse image sequence in under agarose plates coated with HSA showing T_PLN-M_ (top) and T_RM_ (bottom) motility. Cell displacement shown by segmented line. Scale bar, 10 μm. Time in min:s. **F**. Representative T_N_ (n = 75), T_PLN-M_ (n = 226) and T_RM_ (n = 379) tracks. **G.** T_N_, T_PLN-M_ and T_RM_ speeds in under agarose plates coated with HSA. Numbers indicate percentage of tracks > 3 µm/min (boxed). Lines indicate median. **H**. Meandering index of T_N_, T_PLN-M_ and T_RM_ tracks. **I.** T_RM_ speeds after treatment with PTx, RGD peptide, anti-Mac1 mAb, or in presence of EDTA. Numbers indicate percentage of tracks > 3 μm/min (boxed). Lines indicate median. **J.** Image sequence of TRM protrusions in presence of EDTA. Scale bar, 10 µm. **K.** Representative T_RM_ cell tracks in presence of EDTA (n = 75). **L.** Mean displacement over time of T_RM_ tracks. Numbers indicate motility coefficients (μm^2^/min). Data in C, G, H, I and L were pooled from at least 2 independent experiments each. Statistical analysis was performed with unpaired t-test (C) or Kruskal-Wallis with Dunn’s multiple comparison in G - I (as compared to “T_RM_”). **, p < 0.01; ***, p < 0.001.

We then examined T cell displacement on plates coated with fatty acid-free human serum albumin (HSA) and thus free of chemoattractants and specific adhesion ligands. In line with previous findings (*15*), T_N_ and T_PLN-M_ remained essentially immobile throughout the observation period (**Fig. 3, D to F; movies S7 and S8**). Under these conditions, only 16% of T_N_ and 31% of T_PLN-M_ migrated faster than 3 µm/min, and showed low directionality (**Fig. 3, G and H**). In contrast, most SMG T_RM_ showed robust intrinsic motility on HSA-coated plates despite the absence of chemoattractants and adhesion molecules (**Fig. 3, D to F; movies S7 and S8**). Almost 70% of SMG T_RM_ migrated faster than 3 µm/min with high directionality, with their median speed of 5.5 µm/min approaching values observed *in vivo* (**Fig. 3, G and H**). High temporal resolution imaging revealed that migratory T_RM_ often formed several protrusions along the leading edge that appeared to probe the environment, followed by rapid displacement of the cell body along one of the protrusions (**movie S9**). Thus, unexpectedly, confinement alone was sufficient to induce spontaneous SMG T_RM_ migration, representing to the best of our knowledge the first observation of such a motility mode in resting T cells. Their speeds were increased in presence of chemokines and adhesion molecules, suggesting that external promigratory factors tune intrinsic cell motility.

### Friction mediates T_RM_ migration in the absence of chemokines and ICAM-1

We set out to characterize the requirements for autonomous T_RM_ motility in under agarose assays. Reflecting the absence of chemoattractants and integrin ligands, pertussis toxin (PTx) treatment or addition of the β1-blocking peptide RGD did not affect T_RM_ speeds in this setting (**Fig. 3I**). Although Mac-1 binds weakly to serum albumin (*39*), addition of anti-Mac-1 mAb did not cause a significant reduction in T_RM_ speeds (**Fig. 3I**). These observations suggested a friction-based migration mechanism (*40*). Friction is the resisting force when two elements slide against each other and may be composed of a number of fundamental forces. While the nature of the weak interactions between T_RM_ and migratory surface causing friction are not defined, we hypothesized that these might in part involve bivalent cations. Indeed, chelation of bivalent cations by EDTA caused a strong decrease of T_RM_ speeds under agarose (**Fig. 3I**). High temporal resolution imaging showed that despite the lack of translocation in presence of EDTA, T_RM_ continued to probe the environment via transient protrusion formation, essentially “running on the spot” (**Fig. 3, J and K, movie S10**). This behavior precipitated a loss in the motility coefficient (**Fig. 3L**). In sum, our data suggest that bivalent cation-dependent friction between T_RM_ and the confining 2D surfaces generated sufficient traction for translocation in the absence of considerable surface binding.

### SMG T_RM_ insert protrusions between adjacent structures for translocation

In addition to friction-based migration, protrusion insertion has emerged in recent years as a complementary mechanism to allow cell migration without specific adhesions (*40*). The continuous probing of T_RM_ in presence of EDTA (**Fig. 3J**) provided an opportunity to test whether topographic features of the environment such as narrow intercellular spaces may rescue cell motility by permitting insertion of pseudopods as mechanical “footholds” (*41, 42*). As a surrogate approach to re-introduce a “2.5D” environmental geometry in under agarose assays, we co-transferred a surplus of T_N_ together with T_RM_ and performed time-lapse imaging in the presence of EDTA and in the absence of chemoattractants and adhesion molecules (**Fig. 4A**). Remarkably, T_RM_ localized within T_N_ clusters frequently showed lateral displacement despite the presence of EDTA (**movies S10 and S11**). Under these conditions, T_RM_ displacement occurred through insertion of protrusions between adjacent T_N_ and subsequent translocation of the cell body accompanied by dynamic cell shape changes (**Fig. 4B**). T_RM_ within T_N_ clusters were significantly faster than isolated T_RM_ (5.8 ± 3.0 and 2.3 ± 1.7 µm/min, respectively), displayed higher directionality, and resembled in cell shape and speeds T_RM_ migrating *in vivo* (**Fig. 4C to E**). Once T_RM_ had traversed T_N_ clusters, they returned to their probing behavior without efficient translocation, indicating a close interdependence on physical contact and motility (**movie S10**).

**Fig. 4.**
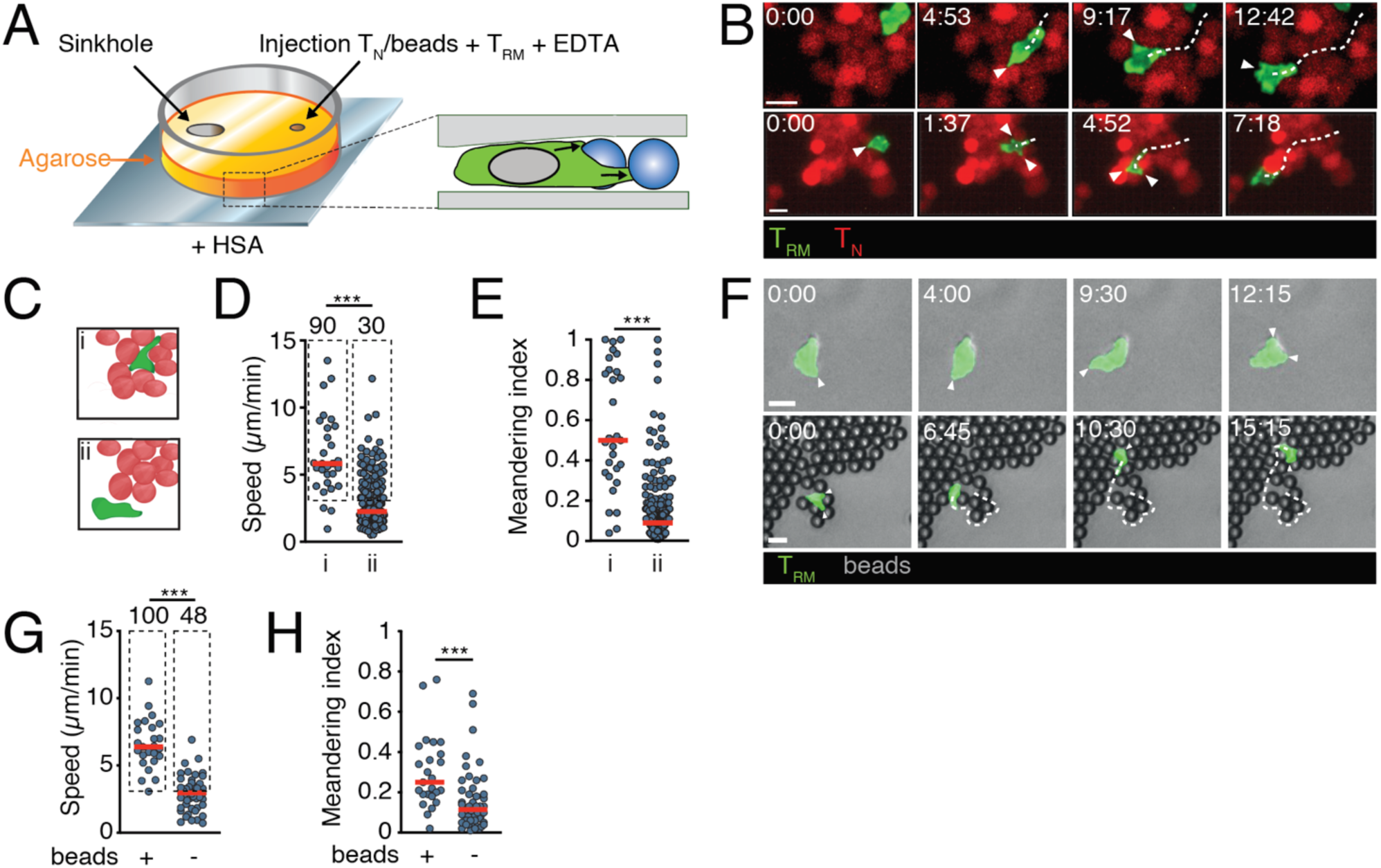
T_RM_ insert protrusions for cell displacement in absence of external chemoattractants and friction. **A.** Experimental layout. Arrows indicate protrusion direction. **B**. Image sequences of T_RM_ within T_N_ clusters in presence of EDTA. Arrowheads show membrane protrusions; segmented line indicates cell track. Scale bar, 10 μm. Time in min:s. **C.** Graphical representation of T_RM_ inside T_N_ cluster (i) or dispersed (ii). **D.** T_RM_ track speeds according to their location. Numbers indicate percentage of tracks > 3 μm/min (boxed). Lines indicate median. **E**. Meandering index of T_RM_ tracks sorted according to their location. Lines indicate median. **F.** Image sequences of T_RM_ alone (top) and with 7 µm polystyrene beads (bottom) in presence of EDTA. Arrowheads show membrane protrusions. Segmented line indicates cell track. Scale bar, 10 µm, time in min:s. **G.** T_RM_ track speeds according to their association with or without beads. Numbers indicate percentage of tracks > 3 μm/min (boxed). Lines indicate median. **H**. Meandering index of T_RM_ tracks sorted according to their location. Lines indicate median. Data in D, E, G and H are pooled from 4 - 5 independent experiments. Statistical analysis was performed with Mann-Whitney test. ***, p < 0.001.

We then examined whether potential residual molecular interactions between naive T cells and T_RM_ might act as drivers of migration. We therefore transferred uncoated polystyrene beads with T_RM_ in under agarose assays. These beads replaced T_N_ as surrogate 2.5D structures and allowed to examine protrusion insertion in the absence of potential adhesive interactions. In this setting, T_RM_ recapitulated the behavior observed within T_N_ clusters, showing effective cell displacement only when in contact with clusters of beads for protrusion insertion (**Fig. 4F; movie S12**). T_RM_ speeds increased to 6.4 ± 1.9 µm/min and became more directional when in contact with beads, whereas isolated T_RM_ showed no displacement (**Fig. 4, G and H**). In this setting, we further observed that T_RM_ moved around dense bead areas, in line with a search for permissive gaps for locomotion (**movie S12**). In sum, SMG T_RM_ displayed a unique ability to migrate by adapting to topographic features of the environment through protrusion insertion and shape deformation, even in the absence of considerable friction, chemoattractants and adhesion receptors.

### Residual in vivo motility of SMG T_RM_ in presence of integrin and chemokine receptor blockade

Our *in vitro* experiments raised the question to which extent external cues govern T_RM_ motility *in vivo*. We explored the molecular mechanisms underlying T_RM_ scanning of SMG, focusing on well-described canonical chemoattractant- and integrin-signaling pathways. Integrins provide traction and force transmission through engagement of their ligands expressed by many cell types including macrophages, such as ICAM-1. SMG T_RM_ express α1, α4, αE, αL, β1, β2 and β7 integrins, and low levels of αV (**Fig. S1 and 4A**). To assess their involvement in SMG T_RM_ immune surveillance, we administered a mix of integrin-blocking mAbs against the major lymphocyte integrin αL (CD11a/CD18, LFA-1), the E-cadherin ligand αE (CD103), and α4 (VLA-4 and α4β7) to LCMV-OVA memory phase mice containing T_PLN-M_ and T_RM_ (**Fig. 5A**). We confirmed that mAbs were saturating surface integrins at the time point analyzed (**Fig. S4B**). We then followed OT-I T cell motility in PLN and SMG on ≥ day 30 p.i., using dual surgery 2PM as described above. Integrin blockade significantly lowered T_PLN-M_ speeds from 11.7 to 8.8 µm/min (**Fig. S4C**), similar to the decreased cell speeds of CD18-deficient T_N_ in lymphoid stroma (*14*). In contrast, T_RM_ speeds and crawling along tissue macrophages remained unaltered by this treatment (**Fig. 5, B and C; movie S13**). We did not detect Mac-1 (CD11b/CD18) expression on SMG T_RM_ by flow cytometry, and addition of anti-Mac1 mAb to the integrin blocking mix did not decrease T_RM_ speeds or guidance by tissue macrophages (not shown). Similarly, inclusion of blocking mAbs against α1 and αV together with αL, α4 and αE had no impact on T_RM_ cells speeds or association with macrophages (6.5 ± 2.7 µm/min; n = 68 tracks).

**Fig. 5.**
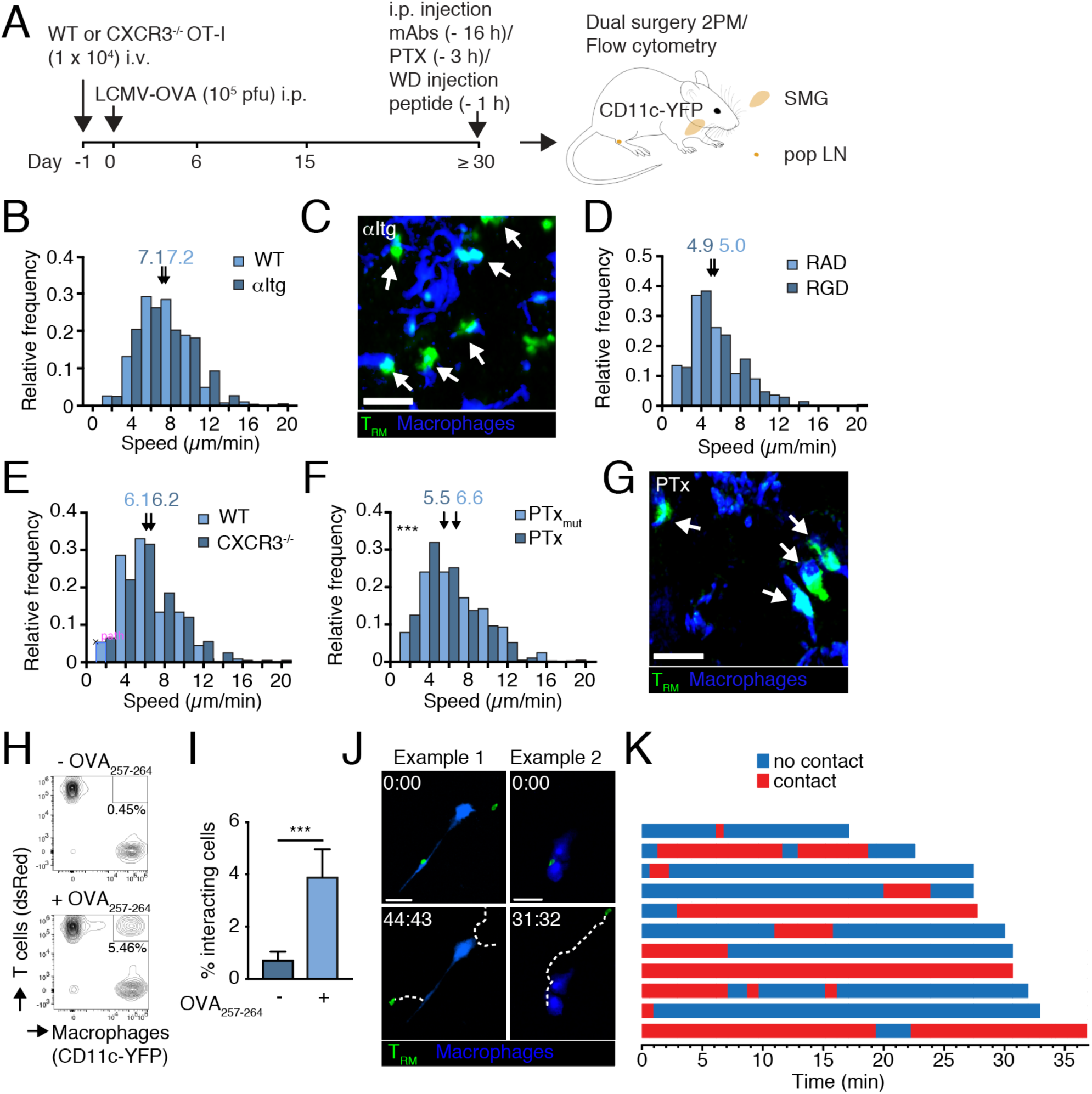
Residual *in vivo* SMG T_RM_ motility during inhibition of Gαi and integrins. **A.** Experimental layout. **B.** OT-I T_PLN-M_ and T_RM_ speeds after combined anti-αL, α4 and αE integrin mAb (αItg) inhibition. Arrows indicate median values (µm/min). **C.** 2PM image of T_RM_ – tissue macrophage colocalization in αItg-treated SMG. Arrows indicate T cell – tissue macrophage contacts. Scale bar, 20 µm. **D.** OT-I T_RM_ speeds in SMG after WD administration of RAD or RGD peptide**. E.** WT and CXCR3^-/-^ OT-I T_RM_ speeds in SMG in memory phase (≥ day 30 p.i.). Arrows indicate median values (µm/min). **F.** OT-I T_PLN-M_ and T_RM_ speeds after systemic treatment with active PTx or inactive (mutant) PTx (PTx_mut_). Arrows indicate median values (µm/min). **G.** 2PM image of T_RM_ – tissue macrophage colocalization in PTx-treated SMG. Arrows indicate T cell – tissue macrophage contacts. Scale bar, 20 µm. **H.** Flow cytometry plot of mixed T_RM_ and macrophages. **I.** Quantification of cluster formation as shown in H. **J.** Example image sequences showing T_RM_ in transient contact with macrophages under agarose on fibronectin-coated plates. T_RM_ displacement is shown by segmented line. Scale bar, 50 μm. Time in min:s. **K.** T_RM_ – macrophage contact duration for individual tracks. Data in B, D, E and F are pooled from 2-5 independent experiments with a total of 2-7 mice with at least 111 tracks per condition and analyzed with unpaired Student’s t-test. Data in I are pooled from 2 independent experiments and analyzed using unpaired Student’s t-test. ***, p < 0.001.

Poor surface saturation of blocking anti-β1 mAbs on OT-I T cells preempted us to assess the role of β1 integrins for T_RM_ motility by this approach (not shown). As alternative, we directly administered the β1-blocking peptide RGD or the control peptide RAD through the Wharton’s duct (WD) into SMG and followed its impact on T_RM_ motility parameters by 2PM. The WD channels saliva from SMG into the oral cavity and can be used to administer reagents or pathogens through retrograde duct cannulation (*43*). Control experiments using WD administration of OVA_257-264_ peptide led to instantaneous arrest of T_RM_ similar to systemic injection, suggesting efficient peptide permeation of SMG by this route (not shown). While WD injection of either peptide slightly lowered T_RM_ speeds, we did not observe any impact on RGD administration on T_RM_ motility parameters as compared to control peptide (**Fig. 5D**). This reflected low β1 integrin levels in the interface between macrophages and T_RM_ (**Fig. S4D**). Furthermore, E-cadherin levels on macrophages and T_RM_ were barely detectable in SMG tissue sections, arguing against a role for this cadherin in mediating close spatial association with tissue macrophages (**Fig. S4E**).

Cytokine-driven chemoattractant production plays a key role for T cell trafficking. Since the CXCR3 ligands CXCL9 and CXCL10 play a role in T_RM_ clustering in SMG (**Fig. 2F**), we co-transferred WT and CXCR3^-/-^ OT-I T cells one day prior to LCMV-OVA infection. Consistent with recent reports (*44*), we found that absence of CXCR3 did not impair T_RM_ formation in SMG after viral infection. Non-clustered CXCR3^-/-^ OT-I T_RM_ showed no significant differences in speeds as compared to WT T_RM_ (**Fig. 5E**), and lack of CXCR3 did not prevent T_RM_ patrolling along tissue macrophages (**Fig. 2F**).

To comprehensively assess a function for potential chemoattractants, we inhibited Gαi signaling by systemic PTx treatment (*45*) and performed 2PM analysis of OT-I T cell motility parameters on ≥ day 30 after LCMV-OVA infection. To control for inhibitor efficacy, we took advantage of the dual surgery of PLN and SMG in the same recipient. Systemic PTx administration significantly slowed T_PLN-M_ down from 11.3 µm/min in control versus 8.6 µm/min in PTx-treated recipients (**Fig. S4F**), resembling observations made with PTx-treated T_N_ in PLN (*11, 13*). Speeds were also decreased in SMG T_RM_ (from 6.6 to 5.5 µm/min) by PTx treatment (**Fig. 5F**), suggesting a role for chemoattractants in mediating high T_RM_ speeds. Nonetheless, we observed a robust residual motility and continued T_RM_ crawling along tissue macrophages in presence of PTx (**Fig. 5G, movie S14**). These data suggest that while Gαi-coupled receptors contribute to SMG T_RM_ motility, they are not required for T_RM_ association with tissue macrophages. Finally, since matrix metalloproteinases (MMP) play roles in cancer cell invasion, we interfered with MMP activity using the broad MMP-9, MMP-1, MMP-2, MMP-14 and MMP-7 inhibitor marimastat as described (*46*). Yet, MMP inhibition did not reduce T_RM_ migration compared to vehicle and rather resulted in a minor increase in speeds (not shown). Taken together, with exception of a minor effect by PTx, the *in vivo* inhibitor treatment examined here did not alter T_RM_ motility and close spatial proximity to tissue macrophages.

To directly assess intercellular adhesion between tissue macrophages and T_RM_ *ex vivo*, we co-incubated freshly isolated cells for 20 min and analyzed cluster formation by flow cytometry (**Fig. 5H**). As positive control for T cell association with tissue macrophages, we pre-incubated macrophages with cognate OVA_257-264_ peptide. While addition of OVA_257-264_ to tissue macrophages induced detectable association with T_RM_, baseline association between both populations remained low (**Fig. 5, H and I**). Finally, we performed under agarose assays in presence of tissue macrophages. On the few occasions when motile T_RM_ contacted co-plated tissue macrophages, these contacts were mostly transient (**Fig. 5, J and K; movie S15**). Furthermore, T_RM_ did not crawl along macrophage protrusions as observed *in vivo* (**Fig. 5J**). These data suggested that T_RM_ association to macrophages occurred preferentially in the SMG microenvironment. Thus, within the technical limitations of our experimental approach, our *in vivo* and *in vitro* observations did not identify specific molecules that provide strong adhesion of SMG T_RM_ to tissue macrophages. Importantly, our data do not exclude the presence of unidentified adhesion receptors mediating T_RM_ association to tissue macrophages *in vivo*.

### Discontinuous macrophage attachment within SMG

We considered that tissue microanatomy may contribute to the close spatial association between T_RM_ and tissue macrophages observed *in vivo*. Based on our observation that *ex vivo* SMG T_RM_ are able to insert protrusions into narrow spaces between adjacent structures lacking adhesion between them (**Fig. 4**), we decided to examine macrophage attachment within SMG applying the super-resolution shadow imaging microscopy (SUSHI) technique. SUSHI was originally developed to visualize the complex topology of the extracellular space (ECS) in living brain slices (*47*). It was used to study dynamic changes in ECS in response to a hyperosmotic challenge, which leads to cell shrinkage and ECS widening in brain tissue. Here, we adapted SUSHI imaging to acutely sliced SMG sections, which were superfused with the cell-impermeable fluorescent dye Calcein (**Fig. 6A**). Steady-state imaging revealed that the interstitium contained more ECS as compared to the tightly packed epithelium (**Fig. 6, B and C**). We reasoned that SUSHI in combination with hyperosmotic challenge could be applied to explore attachment between neighboring cells. Performing time-lapse ECS imaging, we acutely increased the osmolarity to induce cell shrinkage, which led to a strong increase of ECS in the interstitium (**movie S16**). In turn, interepithelial junctions remained relatively stable and only mildly increased their spacing under osmotic challenge, reflecting the presence of adherens and tight junctions known to link epithelial cells (**Fig. 6D**). In contrast, hyperosmolarity induced intraepithelial CD11c-YFP^+^ macrophages detachment from the adjacent epithelium (**Fig. 6, E and F**). This observation confirms previous reports that tissue macrophages do not form continuous adhesive contacts with the epithelium, unlike the extensive cell-to-cell contacts between acinar epithelial cells (*48*).

**Fig. 6.**
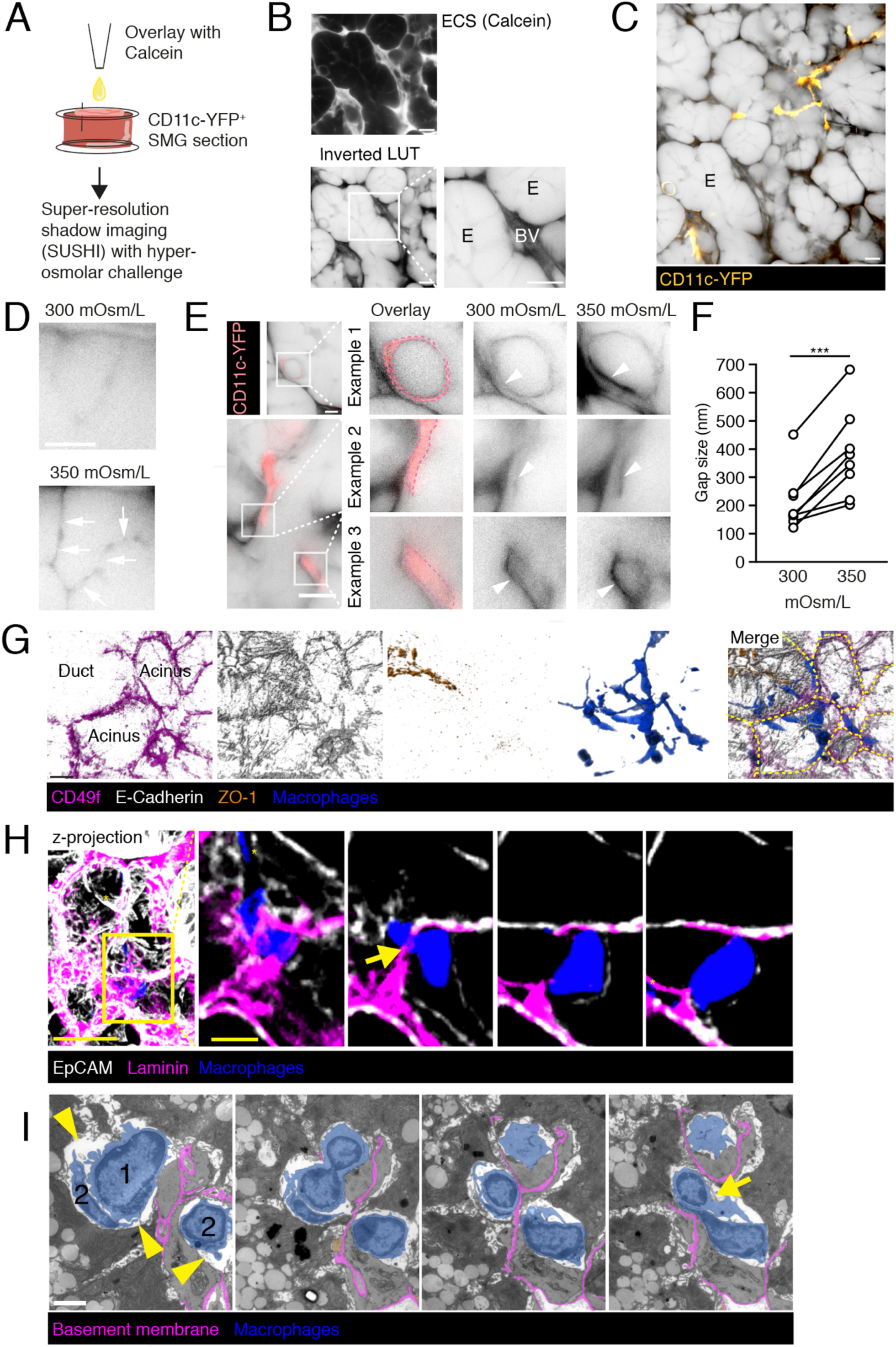
Tissue macrophage attachment in SMG. **A.** Experimental layout of super-resolution shadow imaging (SUSHI) of SMG slices. **B.** Example of SUSHI image for determination of extracellular space. E, epithelium; BV, blood vessel. Scale bar, 10 µm. **C.** Overview of ECS signal with SMG epithelium (E) and CD11c-YFP^+^ tissue macrophages. Scale bar, 10 µm. **D.** Example of epithelial attachment before and after hyperosmotic challenge. Arrows show interepithelial junctions. Scale bar, 5 µm. **E.** Example of macrophage detachment before and after hyperosmotic challenge. Arrowheads indicate detachment. Scale bar, 5 µm. **F.** Quantification of gap size between macrophage and epithelium before and after hyperosmotic challenge. **G.** Immunofluorescent SMG section showing macrophages and epithelial cells in acini and ducts (identified by luminal ZO-1 labelling). Yellow dashed lines indicate outlines of acini and ducts. Scale bar, 10 µm. **H.** Confocal image of SMG section with macrophage protrusions traversing a basement membrane below an epithelial acinus (indicated by arrow). Scale bar, 20 μm (overview) and 5 μm (insert). **I**. Electron microscopy image of macrophages creating a discontinuation of the basement membrane of an acinus (indicated by arrow). Numbers mark two neighboring macrophages. Arrowheads indicate lack of tight adhesion between macrophages and neighboring cells. Scale bar, 2 μm. All images are representative of at least 2 independent experiments. Data in F were analyzed using a paired t-test. ***, p < 0.001.

We then investigated the spatial arrangement of macrophage protrusions with regard to epithelial BM markers. The laminin ligand CD49f (α6) was prominent on the basal side of acini and to a lesser extent on ducts, which were identified by the presence of the tight junction protein ZO-1 on the luminal side. In some cases, tissue macrophages appeared to cross adjacent acini and ducts via their protrusions (**Fig. 6G**). For a detailed examination, we analyzed laminin-stained tissue sections from immunized CD11c-YFP mice. We observed in some cases macrophage protrusions penetrating between adjacent acini, or between epithelium and connective tissue, thus bridging adjacent compartments separated by BM (**Fig. 6H; movie S17**). Using correlative confocal and transmission electron microscopy, we validated that some macrophage protrusions transversed BM (**Fig. 6I**). Taken together, our data support the notion of discontinuous attachment of tissue macrophages to neighboring cells and occasional penetration of macrophage protrusions across the epithelial BM.

### Depletion of tissue macrophages disrupts T_RM_ patrolling

Our observations prompted us to examine T_RM_ motility in the absence of tissue macrophages. To this end, we generated bone marrow chimera by reconstituting C57BL/6 or Ubi-GFP mice with control CD11c-YFP or CD11c-DTR bone marrow. At 6 weeks of reconstitution, we adoptively transferred GFP^+^ or DsRed^+^ OT-I T cells, followed by LCMV-OVA infection. In some experiments, we directly transferred OT-I T cells into CD11c-DTR mice and infected mice with LCMV-OVA. Both approaches allowed us to deplete CD11c^+^ macrophages by diphtheria toxin (DTx) treatment in the memory phase without affecting the unfolding of the adaptive immune response. Macrophage depletion in the memory phase had no impact on CD45^+^ and OT-I T cell numbers recovered from spleens and SMG up to one week after DTx treatment (not shown).

2PM imaging in DTx-treated mice revealed that SMG T_RM_ patrolling behavior was disrupted when macrophages were depleted (**Fig. 7A; movie S18**). T_RM_ motility was decreased, reflected by less displacement (**Fig. 7B**) and slower speeds (**Fig. 7C**). Also, we occasionally observed cells that returned and migrated back the same path within acini and ducts after macrophage depletion (**Fig. 7A**). To quantify this behavior, we developed a method to specifically retrieve U-turns from track parameters (**Fig. 7D**). This analysis confirmed that the percent of T cell tracks showing U-turns was doubled in DTx-treated CD11c-DTR SMG from 8.1 to 16.7 % of tracks (**Fig. 7E**). For comparison, PTx treatment had essentially no impact on U-turn frequency (1.15 fold increase as compared to PTx_mut_). We observed a similar impact of macrophage depletion on T_RM_ speeds in LG (from 7.6 ± 4.3 to 5.5 ± 3.2 µm/min), with a 2.5 fold increase in U-turns (**Fig. S5**).

**Fig. 7.**
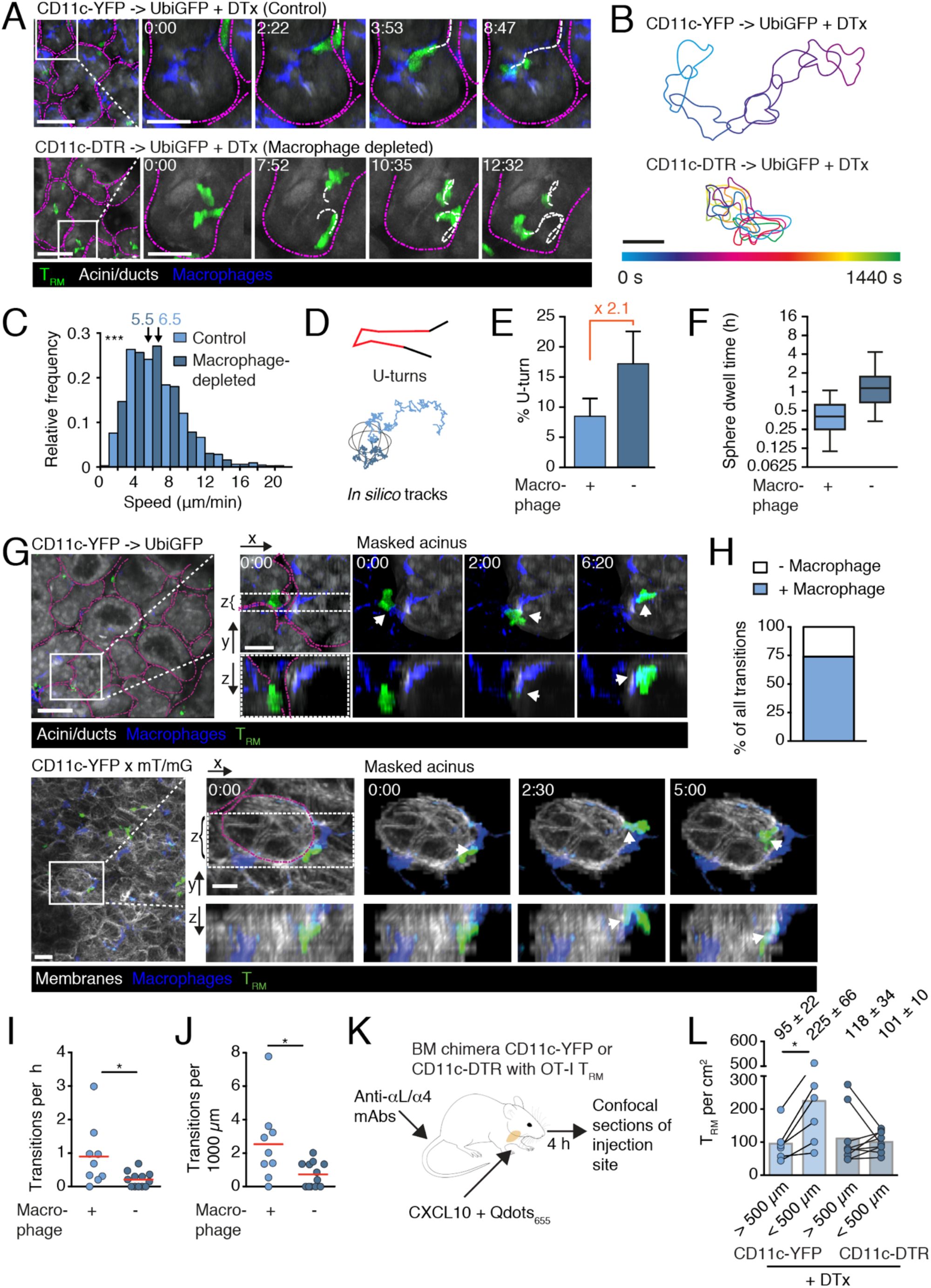
Tissue macrophages assist T_RM_ patrolling of SMG. **A.** 2PM time-lapse image sequence of T_RM_ in DTx-treated CD11c-YFP or CD11c-DTR -> Ubi-GFP chimeras. Magenta lines indicate outlines of acini, white segmented lines indicate cell tracks. Scale bar, 50 µm (overview) and 20 µm (insert). Time in min:s. **B.** Example T_RM_ tracks in presence or absence of macrophage. Scale bar, 10 µm. **C.** Frequency distribution of T_RM_ speeds in DTx-treated CD11c-YFP or CD11c-DTR bone marrow chimera. Arrows indicate median (µm/min). **D.** Track analysis outline. Top panel. U-turns (red) describe tracks reversing direction while excluding continuous turns. Bottom panel. Synthetic tracks were generated to assess dwell time in an 80 µm-diameter sphere (black). One example track is shown for control (light blue) and macrophage-depleted (dark blue) condition. **E.** Percent of tracks making U-turn. Bars indicate 95% confidence intervals. **F.** *In silic*o dwell times for T_RM_ tracks in 80 µm-spheres based on measured track parameters. **G**. 2PM time-lapse image sequences of T_RM_ crawling along a macrophage to enter acini. Epithelial signal was manually masked to show an isolated acinus in zoomed panels. Dashed white line indicates area displayed in xz-view, and arrow indicates T_RM_-macrophage contact. Top: Scale bar, 50 μm (overview) and 20 μm (insert); bottom: Scale bar, 20 μm (overview) and 10 μm (insert). Time in min:s. **H.** Percentage of T_RM_ transitions into or out of acini and ducts in CD11c-YFP -> Ubi-GFP chimeras (n = 42) with and without contact to macrophages. **I, J.** 2PM time-lapse image sequence of CD11cYFP -> Ubi-GFP and DTx-treated CD11cDTR->Ubi-GFP chimeras were analyzed for T_RM_ crossing events (leaving or entering acini). **I** shows average transitions per hour track duration, and **J** depicts transitions per 1000 μm total distance migrated. Data points represent individual image sequences. Line indicates mean. **K.** Experimental layout for analysis of T_RM_ response to local chemokine. CXCL10 was injected with a fluorescent tracer for 4 h to allow T_RM_ accumulation. Integrin blocking mAbs prevent recruitment of circulating T cells. **L.** T_RM_ per cm^2^ at sites of CXCL10 injection in presence or absence of macrophages. Numbers indicate mean ± SD. Data in C, I, J and L are pooled from 2-4 independent experiments with 4-6 mice total. Data in C, I and J were analyzed with Mann-Whitney and data in L were analyzed with Wilcoxon rank test. *, p < 0.05; ***, p < 0.001.

We next asked how impaired motility impacts organ surveillance. We generated tracks *in silico* from the data sets obtained by 2PM imaging of DTx- and control-treated SMG and assessed the average T_RM_ dwell time in a sphere of 80 µm diameter as surrogate epithelial structure (**Fig. 7D**). This analysis uncovered a nearly threefold increased sphere dwell time from 24 ± 1.8 min for control SMG to 69 ± 6.5 min (median ± SEM) for DTx-treated CD11c-DTR SMG (**Fig. 7F**). For comparison, sphere dwell time was increased from 31 ± 1.8 min in PTx_mut_-treated to 46 ± 2.45 min for PTx-treated SMG. Taken together, macrophage depletion disrupted motility parameters and increased the propensity of T_RM_ to make U-turns.

We investigated whether lack of macrophages may also affect T_RM_ transitions into and out of epithelium as part of the impaired motility pattern. To address this point, we developed an approach to optically separate epithelial from connective tissue. We reconstituted irradiated Ubi-GFP mice expressing GFP in all cells with CD11c-YFP bone marrow before transfer of DsRed^+^ OT-I T cells and systemic LCMV-OVA infection. We found that in these chimera, acini and ducts of surgically prepared SMG were GFP^bright^ and readily identifiable by their glandular shapes, whereas connective tissue was GFP^low^. Using case-by-case 3D rendering of 2PM image sequences in memory phase (≥ 30 days p.i. with LCMV-OVA), we observed that DsRed^+^ T_RM_ were not restricted to individual epithelial structures but occasionally crossed between adjacent acini or between epithelial and connective tissue compartments in a bidirectional manner along macrophage protrusions (**Fig. 7G top; movie S19**). We confirmed this observation in a mouse model expressing membrane tomato and CD11c-YFP (**Fig. 7G bottom**). In total, 75% of T_RM_ transits (n = 42) into and out of epithelial structures occurred along macrophage protrusions (**Fig. 7H**). Given that not all tissue macrophages are YFP^+^ (**Fig. S3B**), the actual percentage of macrophage-assisted transitions may still be higher. DTx treatment of CD11c-DTR SMG reduced, but did not abolish, T_RM_ transit into or out of acini and ducts. In total, we observed 55 T_RM_ crossing events into or out of acini in CD11c-YFP versus 12 events in CD11c-DTR chimera SMG. These data corresponded to a 77% fewer crossing events per h track duration and a 71% fewer transitions per 1000 µm track length in macrophage-depleted SMG (**Fig. 7, I and J; movie S18**). Reduced T_RM_ crossing into and out of epithelial structures was also observed when we prolonged DTx treatment for 5 days (**movie S20**).

### Impaired intraorgan accumulation of SMG T_RM_ after macrophage depletion

Tissue macrophages are best characterized for their core functions of maintenance or restoration of tissue homeostasis by engulfing apoptotic cells (efferocytosis), clearing debris and initiation of repair (*49–52*). Accordingly, we observed massively increased numbers of infected cell foci in macrophage-depleted SMG after WD infection with murine cytomegalovirus expressing OVA and mCherry (*53*), as compared to SMG containing tissue macrophages (**Fig. S6, A and B**). The efferocytic function of tissue macrophages was independent of the presence of T_RM_ (**Fig. S6C**), although the latter partially suppressed viral replication as assessed by decreased mCherry intensity in viral foci (**Fig. S6D**). In support of this, we observed CD11c-YFP^+^ cells engulfing MCMV-infected cells after SMG infection (**Fig. S6, E and F**). These observations preempted the use of a viral rechallenge model to assess a function for tissue macrophages in facilitating T_RM_ patrolling.

We therefore designed an alternative experiment to assess the support of SMG macrophages for T_RM_ surveillance and local cluster formation. We treated LCMV-OVA-immunized CD11c-YFP and CD11c-DTR BM chimera mice with DTx, followed one day later by local injection of the CXCR3 ligand CXCL10 into SMG (**Fig. 7K**). We also administered anti-α4 and LFA-1 blocking mAbs that block recruitment of circulating T cells to SMG (*44*) but do not affect T_RM_ motility in this organ (**Fig. 5B**). At 4 h after CXCL10 administration, we isolated SMG and quantified T_RM_ enrichment in thick confocal SMG sections according to the area marked by the co-injected fluorescent marker. CXCR3^-/-^ OT-I T cells did not show accumulation in CXCL10 injection sites, supporting the specificity of chemokine-triggered clustering (49 - 63 > 500 µm versus 60 - 68 cells/cm^2^ cells < 500 µm from injection site; range from two SMG each). WT OT-I T_RM_ were twofold enriched at CXCL10 injection sites, suggesting that these cells had followed a CXCL10 gradient or became retained during their surveillance path (**Fig. 7L**). In contrast, local accumulation of T_RM_ was lost when macrophages had been depleted, although T_RM_ numbers outside the site of chemokine injection remained comparable to macrophage-containing SMG (**Fig. 7L**). These data suggest a key role for SMG macrophages to assist T_RM_ patrolling within and between epithelial structures and to cluster at local inflammatory sites (**Fig. S7**).

## Discussion

After clearing of pathogens, T_RM_ display a remarkable capacity to patrol heterogeneous tissues without impairing vital organ functions (*16–20*). Their scanning behavior evolved because T cells are MHC-restricted and hence need to physically probe membrane surfaces of immotile stromal cells. The key point of this study was to examine how these cells achieve this feat in the complex arborized epithelial structure of SMG during homeostatic immune surveillance. Our main finding is that T_RM_ mostly moved along tissue macrophages, and that depletion of macrophages impaired T_RM_ patrolling. These observations assign a new accessory role to tissue macrophages in addition to their core functions for tissue homeostasis and sentinels of infection. Our data suggest two non-exclusive options to explain macrophage guidance of T_RM_: first, through unidentified specific adhesive interaction(s) independent of ICAM-1 and other adhesion molecules; and second, by offering paths of least resistance within the exocrine gland microenvironment for protrusion insertion by autonomously moving T cells. Our data provide evidence for the second option without discarding the first one. Reductionist *in vitro* experiments revealed that SMG T_RM_ respond to exogeneous cues from chemoattractant and adhesion molecules. Remarkably, confinement alone suffices to trigger friction- and protrusion insertion-based motility without exogeneous chemoattractants or adhesion molecule. We speculate that the continuum of intrinsic motility and integration of external factors permits T_RM_ to patrol these exocrine glands in homeostasis and rapidly respond to inflammatory stimuli.

Macrophages and T cells closely cooperate during the onset of inflammation, the effector phase and contraction through antigen presentation, cytokine secretion and effector functions such as phagocytosis. Yet, little is known whether and how these two cell types collaborate for surveillance of NLT during homeostasis. Tissue macrophages are best characterized for their core function of maintenance or restoration of tissue homeostasis by engulfing apoptotic cells, clearing debris and initiation of repair (*49–52, 54*). A recent study has identified a role for tissue macrophages for cloaking of microlesions (*55*), a behavior we also observed in SMG after local laser injury (not shown). Tissue macrophages also serve as sentinels of infection, leading to cytokine secretion and leukocyte recruitment (*16, 56, 57*). In recent years, several non-phagocytic and non-sentinel functions were assigned to macrophages, as core functions of parenchymal parts of organs are outsourced to accessory cells. Accessory macrophage functions include blood vessel and mammary duct morphogenesis, hematopoietic stem cell maintenance, pancreatic cell specification, lipid metabolism, relay of long-distance signals during zebrafish patterning and electric conduction in the heart (*58, 59*). Our data suggest a novel accessory function, which is to facilitate T_RM_ patrolling within and between acini and ducts of arborized secretory epithelium.

Our initial assumption was that specific adhesion receptors drive T cell association with tissue macrophages, while chemoattractants fuel their high baseline motility. Tissue macrophage express ICAM-1 and other adhesion molecules that can serve as ligands for T cell adhesion receptors, as well as chemoattractants (*38*). It was therefore startling that - against our initial expectations - we were unable to find evidence for strong adhesive contacts between salivary gland macrophages and T_RM_. The experimental systems we have used to address this point encompass *in vivo* inhibition of adhesion receptors in combination with reductionist *in vitro* adhesion assays. Such assays have previously been employed to identify intercellular adhesion through specific molecular interactions, such as ICAM-1-driven binding between T cells and DCs (*60*). It is important to note our data do not rule out the presence of specific adhesive and/or promigratory interactions between T_RM_ and tissue macrophages *in situ*. For instance, low T_RM_ binding to tissue macrophages *in vitro* may be owing to altered gene expression patterns after macrophage isolation (*61*). Along the same line, we have not examined talin-deficient T cells that lack functional integrins, and poor surface mAb saturation preempted an analysis of CD44 for SMG T_RM_ motility (*62*). Of note, PTx treatment induced a minor but significant reduction in T_RM_ speeds *in vivo*. Yet, PTx treatment had essentially no impact on U-turn frequency and movement along tissue macrophages. In line with this, recent observations suggest that guidance and adhesion do not necessarily correlate, as T_N_ migrate along the FRC network even in the absence of LFA-1 and CCR7 (*15*). The influence of the physical properties of the microenvironment is increasingly recognized to play a central role for decision-taking by migrating leukocytes (*63, 64*). Yet, technical limitations in recreating the complex tissue microenvironment of exocrine glands under controlled *in vitro* conditions limit the experimental scope to address this issue in a definite manner.

The canonical model of leukocyte migration postulates chemoattractant-stimulated F-actin polymerization at the leading edge (*4*). The resulting retrograde F-actin flow in turn generates traction and cell body translocation via an integrin “clutch” that binds to adhesion receptors of the ECM or on the surface of neighboring cells. Although integrin-independent migration in 3D matrices has become a widely accepted concept in cell biology based on studies with cell lines and DCs (*40*), several studies uncovered integrin involvement during immune surveillance of skin T cells (*65, 66*). Thus, it remained unclear to which extent integrin-free motility occurs in primary lymphocytes, which contain less cytoplasm and surface area as compared to DCs and cell lines. Another open question was whether memory T cells from distinct anatomical locations would employ similar or tissue-specific mechanisms of host surveillance. Work by Zaid et al. has identified a critical role of G-protein-coupled receptor signaling during scanning by epidermal T_RM_ (*66*). Our own observations confirm that similar to T_N_ and T_PLN-M_, *ex vivo* confined epidermal T_RM_ do not migrate in the absence of integrin ligands or chemoattractants (not shown). Spontaneous motility under 2D confinement appears to constitute therefore a distinctive hallmark of SMG T_RM_ not shared by other resting T cells. Isolated T_RM_ showed high intrinsic protrusive activity *in vitro*, which may reflect high F-actin turnover and/or increased Rho-ROCK-mediated actomyosin contractility. In fact, low adhesiveness under confinement induces spontaneous amoeboid motility via cortical contractility in adherent mesenchymal cell lines (*67, 68*), suggesting that T_RM_ may use a similar mechanism for autonomous migration *in vitro* and *in vivo*. Yet, it remains currently unknown how this unique motility program is imprinted in SNG T_RM_ and whether it is shared by tissue-resident cells from other exocrine glands.

Adhesion-free motility in 2D conditions has been proposed for large, blebbing carcinoma cells, based on friction mediated by a large interface between migrating cells and substrates (*69*). We show that 2D confinement suffices to induce T_RM_ motility through cation-dependent friction, since these cells become unable to translocate their cell bodies in presence of EDTA. Friction is composed of multiple nanoscale forces between two interfaces. For instance, electrostatic and van der Waals forces have been implicated in cell migration and non-specific adherence to substrate (*63, 70, 71*). As chelation of bivalent cations reduced friction below a threshold for cell translocation in our setting, electrostatic forces are likely to be relevant. In principle, cells may compensate for a lower friction by increasing the contacting surface area (*72*). However, lymphocytes are likely too small to generate a sufficiently large interface under these conditions. In turn, T_RM_regained the capability to translocate in presence of EDTA when narrow spaces are created by immotile neighboring cells or beads that lack strong adhesion to each other. This motility mode correlated with continuous changes in cell shapes owing to the intrinsic protrusion formation capacity of T_RM_. Thus, T_RM_ continuously formed multiple simultaneous protrusions that probed the environment, leading to their insertion into permissive gaps and subsequent cell body translocation. How T_RM_ protrusions generated tractive force for cell translocation under these conditions remains incompletely understood. One possibility is that protrusions insert into gaps of the 3D environment akin to cogs of a cogwheel and transmit the necessary force for translocation through retrograde actin flow along irregularly shaped surfaces, even in the absence of adhesion receptors. In fact, this translocation mode is reminiscent of the “squeezing and flowing” mechanism proposed for DCs (*73*), although SMG T_RM_ do not require a chemokine gradient for displacement. We also observed that T_RM_ avoided areas of high bead density, thus choosing the path of least resistance in this mode.

The efferocytic function of tissue macrophages conceivably requires physical contact with surrounding cells to detect and phagocytose senescent or infected cells. Since salivary gland macrophages do not form continuous tight and adherens junctions with neighboring cells (*48*), these cells may create a path of least resistance for patrolling T_RM_. We speculate that the flexible anchorage of macrophage protrusions between epithelial cells may facilitate the insertion of F-actin-rich pseudopods by T_RM_ before squeezing of the nucleus as biggest organelle (*40, 42, 74*). T_RM_ migration along macrophages may be further assisted by unknown adhesion receptors or other molecular interactions between these cells. In any event, the non-proteolytic path finding is beneficial to preserve the integrity of the target tissue, as it does not require constant repair of newly generated discontinuities in the ECM (*75*). The scanning strategy adopted by T_RM_ resembles the migration pattern of T cell blasts in 3D collagen networks, where these cells routinely bypass dense collagen areas, while probing the environment for permissive gaps for cell body translocation (*76*). In fact, leukocytes have recently been shown to use the nucleus to identify the path of least resistance in complex 3D environments with different pore sizes (*64*). This migration mode preserves tissue integrity is energetically favorable by avoiding ECM degradation.

Reflecting the multiple functions of tissue macrophages, depletion studies make the interpretation of the physiological function of macrophage-assisted T_RM_ surveillance of SMG experimentally difficult to dissect. As example, when we locally infected macrophage-depleted SMG with MCMV, we observed massively increased numbers of viral foci as compared to control SMG owing to a lack of efferocytosis. Our local CXCL10 deposition experiment in combination with impaired recruitment of circulating T cells suggests that macrophages facilitate local T_RM_ accumulation at sites where inflammatory chemokines are produced. This resembles observations made in skin infection models where CXCR3 promotes CD8^+^ T cell accumulation at sites of viral replication necessary for efficient elimination of infected cells (*21, 77*). A recent study by Förster and colleagues has uncovered an unexpectedly low killing rate of cytotoxic T cells against viral-infected stromal cells (*78*). Thus, effective stromal cell elimination requires cooperativity through repeated cytotoxic attacks by multiple CD8^+^ T cells. Conceivably, the promigratory accessory function of tissue macrophages described here helps to cluster a quorum of T_RM_ for successful stromal cell killing. Furthermore, unlike the monoclonal T_RM_ population created in our experimental setting, not all T_RM_ recognize the same pathogen under physiological conditions. This might impose a requirement for T cells to scan local sites of pathogen re-emergence and to form clusters for timely elimination of fast-replicating microbes.

In sum, our data assign a previously unnoticed interplay between tissue-resident innate and adaptive immune cell populations. These findings further suggest a noticeable capacity of SMG T_RM_ to integrate a continuum of intrinsic and external signals, friction and 3D structures for efficient motility, providing these cells with maximal flexibility for NLT surveillance. We propose that such a mode of tissue patrolling is ideally adapted to the arborized epithelial architecture of exocrine glands by permitting homeostatic surveillance while maintaining responsiveness to local inflammatory cues.

## Materials and Methods

### Mice

OT-I TCR (*34*) and P14 TCR transgenic mice (*36*) were backcrossed to Tg(UBC-GFP)30Scha “Ubi-GFP” (*79*) or hCD2-dsRed (*80*) mice. Ubi-GFP (GFP^+^) OT-I mice backcrossed to CXCR3^-/-^ mice have been described (*81*). Tg(Itgax-Venus)1Mnz CD11c-YFP (*82*) and Tg(Itgax-DTR/EGFP)57Lan CD11c-DTR mice were used as recipients or bone marrow donors for lethally irradiated C57BL/6 or Ubi-GFP mice. C57BL/6 mice were purchased from Janvier (AD Horst). All mice were maintained at the Department of Clinical Research animal facility of the University of Bern, at the Theodor Kocher Institute and the University of Fribourg. All animal work has been approved by the Cantonal Committees for Animal Experimentation and conducted according to federal guidelines.

### T cell transfer and viral infections

CD8^+^ T cells were negatively isolated from spleen, peripheral and mesenteric lymph nodes of GFP^+^ or dsRed^+^ OT-I or GFP^+^ P14 mice, using the EasySep^TM^ Mouse CD8^+^ T cell Isolation Kit (Stem Cell Technologies). CD8^+^ T cell purity was confirmed to be > 95% by flow cytometry prior to cell transfer. 10^4^ OT-I T cells were i.v. transferred into recipient mice 24 h before i.p. infection with 10^5^ pfu LCMV-OVA (*33*). Experimental read-outs for the acute, cleared and memory phase of viral infection were performed 6, 15 and ≥ 30 days p.i., respectively.

### LCMV virus titer

C57BL/6 mice were infected i.p. with 10^5^ pfu LCMV-OVA and sacrificed 3 or 5 days later. PLN, spleens and SMG were harvested and organs were snap frozen in liquid nitrogen. Recombinant LCMV-OVA infectivity was measured by immunofocus assay on MC57 cells as previously described ^96^.

### Antibodies and reagents

Alexa633-conjugated anti-PNAd MECA79, αL-integrin FD441.8 and anti-α4-integrin PS/2 mAbs were from nanotools (Freiburg, Germany). Anti-α1-integrin Ha31/8 was from BD Bioscience, anti-α4 integrin PS/2 and anti-αE integrin M290 were from BioXCell, anti-αV integrin RMV-7 was from BioLegend, and anti-Mac1 mAb M1/70 was purified from hybridoma supernatant. TexasRed-Dextran 70 kDa was from Molecular Probes. Cascade Blue (MW 10 kDA) was purchased from Invitrogen. TRITC-Dextran (MW 70 kDa) and Diphtheria Toxin whereas purchased from Sigma. Pertussis toxin (PTx) and enzymatically inactive mutant PTx (PTx_mut_) were obtained from List Biological Laboratories. Sodium Pyruvate (100 mM; #11360-039), HEPES buffer (1M; #15630-056), Minimum essential Medium Non-essential amino acids (MEM NEAA, #11140-035), L-Glutamine (200 mM; #25030-024), PenStrep (#15140-122) and RPMI-1640 (#21875-034) were purchased from Gibco and Fetal Bovine Serum (FCS, #SV30143.03) was purchased from HyClone.

### Flow cytometry analysis

PLN and spleen were harvested at the indicated time points and single cell suspensions were obtained by passing organs through cell strainers (70 µm; Bioswisstec). Red blood cell lysis was performed on splenocytes in some experiments. For analysis of SMG and LG, organs were minced and treated with 2 U/µl collagenase II (Worthington Biochem), 2 U/µl bovine DNAse I (Calbiochem) and - only for intracellular stainings of cytokines – 5 µg/ml Brefeldin A (B6542, Sigma-Aldrich) in CMR (RPMI/10% FCS/1% HEPES/1% PenStrep/2 mM L-Glutamine/1 mM Sodium Pyruvate) for 30 min at 37°C, passed through a 70 µm cell strainer and washed with PBS/5 mM EDTA. We used following reagents for flow cytometry:

| Antibody | clone | company | Order number |
| --- | --- | --- | --- |
| anti-CD3-APC | 145-2C11 | Biolegend | 100312 |
| anti-CD8a-PE | 53-6.7 | BD Biosciences | 553033 |
| anti-CD8a-PerCP | 53-6.7 | Biolegend | 100732 |
| anti-CD8a-APC/Fire750 | 53-6.7 | Biolegend | 100766 |
| anti-CD11a-PE | M17/4 | Biolegend | 101107 |
| anti-CD11b-PE | M1/70 | BD Biosciences | 553311 |
| anti-CD11c-APC | HL3 | BD Biosciences | 550261 |
| anti-CD18-PE | M18/2 | Biolegend | 101407 |
| anti-CD29-PE | HMβ1-1 | Biolegend | 102207 |
| anti-CD44-PE | IM7 | BD Biosciences | 553134 |
| anti-CD45-PerCP | 30-F11 | BD Bioscience | 557235 |
| anti-CD45-BV711 | 30-F11 | Biolegend | 103147 |
| anti-CD45.1-AF488 | A20 | Biolegend | 110718 |
| anti-CD45R/B220-APC | RA3-6B2 | Biolegend | 103212 |
| anti-CD49a-PE | HMα1 | Biolegend | 142603 |
| anti-CD49b-Biotin | DX5 | Biolegend | 108903 |
| anti-CD49d-PE | PS/2 | Southern Biotech | 1520-09L |
| anti-CD51-PE | RMV-7 | Biolegend | 104105 |
| anti-CD64-AF647 | X54-517 | BD Bioscience | 558539 |
| anti-CD69-PE | H1.2F3 | Biolegend | 104508 |
| anti-CD103-APC | 2E7 | Biolegend | 121414 |
| anti-CD103-Biotin | M290 | BD Bioscience | 557493 |
| anti-β7-integrin-Biotin | FIB504 | Biolegend | 321209 |
| anti-F4/80-FITC | BM8 | Biolegend | 123108 |
| anti-F4/80-APC | BM8 | Biolegend | 123116 |
| anti-KLRG1-PE | 2F1 | BD Bioscience | 561621 |
| anti-KLRG-1-PE-Cy7 | 2F1 | Biolegend | 138415 |
| anti-KLRG1-APC | 2F1 | Biolegend | 138411 |
| anti-NK1.1-APC | PK136 | Biolegend | 108710 |
| anti-Siglec-F-PE | E50-2440 | BD Bioscience | 562068 |
| Streptavidin-APC | - | Biolegend | 405207 |
| anti-rat IgG1 K-APC | - | Biolegend | 400412 |
| armenian hamster IgG-PE | - | Biolegend | 400907 |

Single cell suspensions were stained for surface antigens on ice for 30 min with the indicated antibodies and washed in FACS buffer (FB; PBS/2% FCS/1 mM EDTA) or FB with 5 µg/ml Brefeldin A for intracellular cytokine stainings. All sample were washed in FB after staining, and for intracellular stainings, cells were permeabilized and fixed in Cytofix/Cytoperm (#51-2090KZ, BD Biosciences) for 20 min on ice. Fixative was removed by washing with Perm/Wash buffer (#51-2091KZ, BD Biosciences) and subsequent intracellular staining steps were performed in Perm/Wash buffer. Cells were washed again prior to acquisition and at least 10^5^ cells in the lymphocyte FSC/SSC gate were acquired using a FACSCalibur (BD Bioscience), LSR II (BD Bioscience), LSR II SORP Upgrade (BD Bioscience) or Attune NxT Flow cytometer (ThermoFisher). Total cell counts were obtained by measuring single cell suspensions in PKH26 reference microbeads (Sigma) for 1 min at high speed. Gating for CD103^+^ and KLRG1^+^ was set according to isotype controls. For CD69 staining, positive and negative gates were set according to distinguishable populations and FMO was subtracted from the final % of CD69^+^ cells as background.

### Immunofluorescence

Mice were anesthetized with i.p. injection of ketamine and xylazine and perfused with ice-cold 1% PFA. Organs were harvested and fixed overnight in 2% PFA prior to embedding in TissueTek O.C.T. compound (Sakura) for cryostat sectioning or 5% low-melting-point agarose (Sigma) for vibratome (Microslicer™ DTK-1000) sectioning. 6 µm-thick frozen cryostat sections were permeabilized, blocked and stained with 0.05% Triton-X 100 in 5% skimmed milk or 0.05% Tween 20 and 3% BSA for 1h, washed 3 times with PBS/1% BSA/0.05% Tween (TBPBS) and stained with goat-anti-Iba1 1/200 (ab5076, Abcam) and anti-phosphotyrosine (pTyr) (ab179530, Abcam) for 2 h at RT prior to mounting with Fluoromount-G (Electron Microscopy Sciences).

For vibratome sections, 100 µm-thick section were collected in a 48-well plate and blocked with TBPBS for 2 h, then blocked with F_C_-block o.n. at 4°C (hybridoma supernatant; 2.4 mg/ml diluted 1/800 in TBPBS). After washing once with TBPBS for 1 h, sections were stained in TBPBS for 2-3 days at 4°C (in 100 µl, 3 sections per well) with Alexa647-conjugated anti-EpCAM (1/160 dilution; clone G8.8, 118212, Biolegend), eFluor660-conjugated anti-E-cadherin (1/200 dilution; clone DECMA-1, 50-3249-1633, eBioscience), polyclonal rabbit anti-Laminin (1/1000 dilution; Z0097, Dako) or Cy3-conjugated anti-α-smooth muscle cell actin (clone 1A4, C6198, Sigma). Sections were washed 3 times for 1 h with TBPBS and incubated with secondary Cy3-conjugated anti-rabbit Ig (1/400 in TBPBS; 111-165-144, Jackson Immune Research), then washed 3 times 1 h with TBPBS and one time with PBS. Images were acquired with a Zeiss LSM510 or Leica SP5 confocal microscope and processed using Adobe Photoshop CS6 and Imaris 8.4.1. We used Imaris software for surface rendering and channel masking function to separate fluorophores with close emission spectra (i.e. GFP and YFP).

### 2PM image acquisition and analysis

2PM intravital imaging of the popliteal lymph node was performed as described (*83*). In brief, mice were anesthetized with ketamine/xylazine/acepromazine. The right popliteal lymph node was surgically exposed. Prior to recording, Alexa 633-conjugated MECA-79 (10 µg/mouse) was injected i.v. to label HEV. 2PM was performed with an Olympus BX50WI microscope equipped with a 20X Olympus (NA 0.95) or 25X Nikon (NA 1.0) objective and a TrimScope 2PM system controlled by ImSpector software (LaVisionBiotec). Some of the image series were acquired using an automated system for real-time correction of tissue drift (*84*). For 2-photon excitation, a Ti:sapphire laser (Mai Tai HP) was tuned to 780 or 840 nm. For 4-dimensional analysis of cell migration, 11 to 20 x-y sections with z-spacing of 2-4 µm (22-64 µm depth) were acquired every 20 s for 20-60 min; the field of view was 150-350 x 150-350 µm. Emitted light and second harmonic signals were detected through 447/55-nm, 525/50-nm, 593/40-nm and 655/40-nm bandpass filters with non-descanned detectors in case of C57BL/6 recipient mice. For CD11c-YFP^+^ recipient mice or bone marrow chimera, we used 447/55-nm, 513/20-nm, 543/30-nm and 624/30-nm bandpass filters.

For imaging of the SMG, neck and thorax of the mouse were shaved, and residual hair removed with hair removal cream (Veet). Subsequently, the animal was fixed on its back onto a custom-built SMG imaging stage and stereotactic holders were attached to the head for stabilization. A 10 x 5 mm piece of skin on the right side of the neck was excised to expose the right SMG lobe, which was micro-surgically loosened from surrounding tissue. The right SMG lobe was flipped to the right and gently immobilized in between 2 cover glasses to minimize motion artifacts from heartbeat and breathing. During the whole operation and imaging procedure tissue was kept moist. During imaging, the temperature at the SMG was monitored and kept at 37°C by a heating ring. In most experiments, mice were operated twice (for PLN and SMG) in alternating order to directly compare behavior of cells in different organs of the same recipient. Prior to imaging, blood vessels were labeled by i.v. injection of 400 – 600 µg of 10 kDa Cascade-blue dextran or 70 kDa TexasRed Dextran. Surgical exposure of the LG was essentially performed as for the SMG, with the mouse fixed on its left flank onto the custom-built SMG imaging stage and a 10 x 5 mm piece of skin excised between the right ear and eye of the mouse.

Sequences of image stacks were transformed into volume-rendered four-dimensional videos with Volocity 6.0 or Imaris 6.00-9.00 (Bitplane), which was also used for semi-automated tracking of cell motility in three dimensions. Drift in image sequences was corrected using a MATLAB script recognizing 3D movement in a reference channel or by using the correct drift function of Imaris. Since our filter set up does not allow complete separation of GFP and YFP signals, we performed spectral unmixing of GFP and YFP using the Image J plugin “Spectral_Unmixing” from Joachim Walter. Cellular motility parameters were calculated from x, y, and z coordinates of cell centroids using Volocity, Imaris and MATLAB protocols. The motility coefficient, a measure of the ability of a cell to move away from its starting position, was calculated from the gradient of a graph of mean displacement against the square root of time. We defined U-turns as the steepest turn over five steps of a track, if it is over more than 166 degrees and has a skew line distance between the first and last step smaller than one mean step of the respective track (**Fig. 7D**) to exclude continuous turns. The given binomial proportion 95% confidence intervals are Wilson Score intervals. We generated 100 synthetic tracks of 12 h duration for each condition using a sampling strategy, which was designed to preserve the correlation between velocity and turning angle and the autocorrelation of velocity and turning angle (*10*). We then took the first timestep further than 40 µm away from the origin of each track as simulated dwelling time in an acinus of 80 µm diameter. These analyses were performed using scientific computing packages for Python. For the image series depicted in the Figures, raw 2PM data was filtered with a fine median filter (3×3×1), and brightness and contrast were adjusted. Shape factors were determined by rendering and tracking cells in Imaris, and manually excluding all cells that did not move along a horizontal axis. The signal from the filter cells was projected into a single z-slice and the shape-factor of the 2D image calculated with Volocity.

### In vivo inhibitor treatment

Gαi signaling by chemokines was blocked as described previously (*45*). Briefly, mice were treated with 3 µg PTx or PTx_mut_ by i.p. injection 3 h prior to imaging. For depletion of CD11c-positive cells in CD11c-DTR mice or BM chimera, 4 ng/g Diphtheria Toxin (DTx) was i.p. injected 24 h prior to imaging. Depletion efficiency of CD11c^+^ cells was determined by flow cytometry and was above 98% in all organs analyzed. For the synchronous blocking of integrins, 100 µg each of the purified mAbs M290, FD441.8, M1/70 and PS/2 were injected i.v. 16 h prior to imaging. Surface saturation of blocking mAbs in PLN and SMG suspensions was determined at the end of the experiment by sample staining with or without the same mAb clones used for blocking, followed by a fluorescently labeled secondary mAb and flow cytometry. RGD peptide or as control GRADSP (RAD) peptide (SIGMA) was injected via Wharton’s duct cannulation (approximately 600 nmol of either peptide in 30 µl per lobe in PBS), as described previously (*43*). For this procedure, mice were anesthetized with ketamine/xylazine and their upper incisors rested on a metal rod and the lower incisors pulled down with string, which kept the mouth open. With the aid of a stereomicroscope, we located the orifice of the Wharton’s duct in the sublingual caruncle and inserted a pointed glass-capillary (Untreated Fused Silica Tubing - L × I.D. 3 m × 0.10 mm, #25715, Sigma). The glass capillary was connected to a Hamilton Micro-syringe (Hamilton) via fine bore polythene tubing (0.28 mm, #800/100/100, Smiths), which allowed the injection of small volumes. For inhibition of MMPs, Marimastat (#S7156) was obtained from Sellcheck and diluted in PBS/10% DMSO (0.2 mg/g) or the corresponding volume of PBS/10% DMSO was injected i.p. 90 min before starting imaging (*46*). OVA_257-264_ (#BAP-201) and gp_33-41_ (#BAP-206) peptides were obtained from ECM microcollections and 200 μg/100 µl saline injected i.v. immediately prior to imaging or 6 - 12 h prior to organ harvest for FACS staining.

### Viral infection via Wharton’s duct cannulation

Wharton’s duct cannulation was prepared as described above. Approximately 12500 pfu MCMV-OVAmCherry (*85*) were injected into the Wharton’s duct (WD) of DTx-treated CD11c-DTR or CD11c-YFP mice in memory phase of LCMV-OVA infection. Mice were euthanized 48 h post infection and SMG tissue fixed in 4% PFA at 4°C for 12 h.

### Under agarose assays

T_N_ were isolated from spleen and PLN of a naive mouse using CD8^+^ T cell isolation kit from Stemcell. SMG-derived macrophages were isolated from uninfected CD11c-YFP mice and sorted for CD11c-YFP^+^ cells. T_RM_ and T_CM_ were isolated from SMG and PLN respectively of > 30 d LCMV-OVA-infected C57BL/6 mice. Single cells suspension of SMG and PLN were stained with APC-conjugated anti-KLRG1 mAb and sorted for GFP^+^ or DsRED^+^ KLRG1^-^ T_RM_ and T_PLN-M_, respectively. A 17-mm diameter circle was cut into the center of 60-mm dishes. The hole was sealed from the bottom part of the dish using aquarium silicone (Marina) and a 24-mm glass coverslip. After the silicone dried, we overlaid a 5 mm-high ring cut from a 15-ml falcon tube and sealed the borders with low melting point paraffin. Coverslips were washed with PBS and coated with 3% human serum albumin (HSA; A1653, Sigma) o.n. at 4°C or for 3 h at 37°C. In some experiments, coverslips were coated with 10 μg/ml fibronectin (11080938001, Roche315-02, PeproTech). Fresh medium was added every 2 days and macrophages were cultured for 6-7 days. For naïve T cell migration, coverslips were coated with 20 µg/ml Protein A (6500-10, BioVision) for 1 h at 37°C, washed 3 times with PBS and blocked with 1.5 % BSA for 2 h at 37°C or o.n. at 4°C. After washing once with PBS, cover glasses were coated for 2 h at 37°C with 100 nM recombinant ICAM1-F_C_ (796IC, R&D Systems) and washed 2 times with PBS. Five ml of 2 x HBSS and 10 ml of 2 x RPMI containing 1% HSA for T_RM_ and T_CM_ and 20% FBS for macrophage and naïve T cell experiments, were mixed and heated in a water bath to 56°C. Golden agarose (100 mg; 50152, Lonza) was dissolved and heated in 5 ml distilled water before adding to the prewarmed medium to give a 1% agarose mix. After cooling to 37°C, 500 µl of the agarose mix was added on top of the coverslip. In some cases, inhibitors were added (200 µg/ml PTx, 5 µg/ml anti-Mac1 mAb, 10 µM RGD or GRADSP, 2.5 mM EDTA, or 5 µg/ml Hoechst (H21492, Invitrogen). After incubation for 30 min at 4°C, the dish was warmed up to 37°C before adding 1 ml of PBS outside the ring to prevent agarose drying. We punched a sink hole (diameter approximately 2 mm) at the side of the agarose. Sorted T cell populations were suspended in RPMI/0.5% HSA and in some cases treated with 5 µg anti-Mac1, 10 µM RGD or GRADSP, and pelleted in an Eppendorf tube. Cells were resuspended in the smallest possible achievable volume (ca. 5-10 µl) and 0.3 µl were injected in the opposite side from the sink hole using a 2.5-µl Eppendorf pipette. In some experiments, polystyrene beads (Sigma-Aldrich, LB30 or 78462) were co-injected with the cells. From the sink hole surplus of medium was collected to confine cells between the agarose and the glass slide. Time-lapse images were taken from the center of the dish using a Zeiss fluorescent microscope (AxioObserver, Zeiss).

### Correlative Confocal and Transmission Electron Microscopy

Correlative confocal and transmission electron microscopy (TEM) was carried out as described (*86*). Briefly, CD11c-YFP mice were perfused with PBS and SMG were fixed *in situ* by left ventricle injection of 1.5% glutaraldehyde/2% PFA in 0.1 M sodium cacodylate buffer (pH 7.4). SMG were harvested and immersed in the same solution for 16 h. Fixed samples were cryoprotected in 30% sucrose prior to embedding in OCT and freezing. Thirty μm sections were cut with a CM1520 cryostat (Leica) and collected on Superfrost Plus slides (Thermo Fisher Scientific). Sections were processed for confocal imaging using PBS as mounting medium to prevent dehydration. After confocal image acquisition, the coverslips were gently removed and sections adherent to the slide were processed for TEM as described (*86*). Briefly, sections were postfixed using the ferrocyanide-reduced osmium-thiocarbohydrazide-osmium (R-OTO) procedure, *en bloc* stained in 1% uranyl acetate and dehydrated through increasing concentration of ethanol. Finally, sections were embedded by overlaying a BEEM capsule filled with Epoxy resin. The BEEM capsules containing the embedded sections were detached by immerging the slides in liquid nitrogen, leaving the section facing up on the resin block. The specimens were mounted on a Leica Ultracut UCT and 70-90 nm thick serial sections were collected on formvar-coated copper slot grids and imaged with a ZEISS Leo912AB Omega fitted with a 2k × 2k bottom-mounted slow-scan Proscan camera controlled by the EsivisionPro 3.2 software. Using the florescent confocal and bright field images, the same areas were relocated in the electron microscope and several images were acquired through the different serial section. Acquired TEM images were then aligned and overlaid with the confocal images by means of the eC-CLEM Icy plugin (*87*).

### Super-resolution shadow imaging (SUSHI)

After euthanizing mice with CO_2_, submandibular salivary glands (SMG) were isolated from 8-11 weeks old C57BL6 or CD11c-YFP mice and submerged in ice-cold PBS. SMG were embedded in 4% low gelling agarose (Sigma), cut in 300 μm-thick transversal slices and submerged in cold complete RPMI medium containing: 10% FCS (Hyclone). Slices were left to recover at room temperature for 15-30 min before entering the imaging chamber of a custom-built 3D-STED microscopy setup (*47*). First, the positively labelled (YFP) macrophages were identified at a depth of 20-30 μm below the surface and imaged in STED mode (excitation 485 nm, depletion 597 nm, objective HC PL APO 63X/1.30 NA, Leica) with the following acquisition parameters: field of view: 200 μm X 200 μm; pixel size: 48 nm X 48 nm; pixel dwell time: 30 μs, frame acquisition time: 20 min. The medium was exchanged in the chamber with the complete RPMI containing 400 μM Calcein dye, which was allowed for 20 - 30 min to disperse throughout the extracellular space of the tissue. Subsequently, we acquired a SUSHI image to identify a region of interest around the macrophages. We performed a hyperosmolar challenge by exchanging the chamber solution (300 μl) with high osmolar solution (350 mOsm/L), and acquired time lapse images to track changes in ECS topology with a 20-min interval between the image frames. All image analysis including morphological measurements were done on raw images using the “Plot Line Profile” function in ImageJ on structures of interest. Brightness and contrast were adjusted using the “Brightness and Contrast” function in ImageJ. It was applied for illustration purposes only and did not affect the quantitative analysis. No filtering or any other image processing was applied, other than inverting the look-up-tables (LUT). The YFP signal was used only to identify macrophages and was not recorded during subsequent imaging. Only unambiguously recognizable macrophages and ECS were analyzed. Image analysis was performed on SMG slices from two mice in two independent experiments.

### Confocal imaging of the p-Tyr signal and quantification

Mice were perfused with PBS containing 4% paraformaldehyde, SMG were isolated and fixed in the same solution at 4°C for 18 h, followed by at least 5 h dehydration in 30% sucrose. Glands were embedded in OCT (Tissue-Tek) and cut at a thickness of 6 and 20 μm at the cryostat, flash dried and fixed with 4% PFA for 10 min at room temperature. Sections were permeabilized using 0.2% Triton X-100, blocked in 10% serum of the secondary antibody and 2% BSA containing PBS and stained with antibodies for 18 h at 4°C in the same solution, after being washed in PBS and mounted in Prolong Gold containing DAPI (Invitrogen, Carlsbad, CA). Fluorescence microscopy was performed using the LSM880 confocal microscopes with 40x oil (Plan-Apochromat 40x 1.3 Oil DIC M27) or 63x oil (Plan-Apochromat 63x 1.3 Oil DIC M27) objectives (Zeiss, Oberkochen, Germany). All images were recorded using sequential excitation. The lack of spectral overlap was confirmed using single fluorescing specimens and antibody specificity via secondary controls. Macrophages were identified via iba-1 and the presence and location of p-Tyr signal was quantified in 7 fields of view using Imaris software. Brightness and contrast were adjusted for each image individually. Gaussian filters were applied using Imaris software.

### Chemokine-driven T_RM_ accumulation

After 6 weeks of reconstitution with CD11c-YFP or CD11c-DTR BM, we transferred 10^4^ GFP^+^ OT-I T cells and infected the day after with 10^5^ pfu LCMV-OVA. After ≥ 30 days p.i., mice were anesthetized one day after i.p. injection of DTx (4 ng/g) as for 2PM imaging. To block immigration of cells from blood, we treated mice with integrin blocking antibodies anti-αL (FD441.8) and anti-α4 (PS/2) (each at 50 µg/mouse; nanotools). SMG was surgically exposed. Using thin glass capillaries as for Wharton’s duct injection, we injected 2 µl of a 1:1 mix of mCXLC10 (100 µg/ml; R&D 466-CR-010) and Qdots_655_ (0.16 µM; Thermo Fisher Q2152MP), to obtain a final mCXCL10 amount of 0.5 µg per site of injection. After 4 h, we sacrificed the mice and harvest the SMG for vibratome sectioning. Mosaic images were taken of 100 µm-thick sections, and lobes with the highest Qdot signal were analyzed by transforming the 3D image into extended 2D image, using the Z-projection function of ImageJ. T_RM_ density in the surrounding area and injection area (defined as an octagon with 500 µm diameter) was calculated using Imaris 8.4.1.

### Statistical analysis

Two-tailed, unpaired Student’s t-test, Mann-Whitney U-test, one-way ANOVA with Dunnett’s multiple comparisons test, Kruskal-Wallis test, or a Wilcoxon rang test was used to determine statistical significance (Prism, GraphPad). Significance was set at p < 0.05.

## Acknowledgements

We thank Dr. Marcus Thelen (IRB, Bellinzona) for support with confocal imaging. This work benefitted from optical setups of the Microscopy Imaging Center of the University of Bern and of the BioImaging platform of the University of Fribourg.

## Author contributions

BS, FT and XF performed most experiments with support by LMA and NR. LMA and KI carried out SUSHI imaging under supervision of UVN. PG carried out computational analysis under supervision of JS. AR and FM performed correlative electron microscopy of SMG sections under supervision of MI. NP, KAK, FB, DM and OTF provided vital material and support. SMSF, MSD and CS analyzed human SMG sections. BS, FT, XF and JVS designed experiments and wrote the manuscript with input from all coauthors.

