## Supplemental_Figures for "Salivary gland macrophages assist tissue-resident CD8^+^ T cell immune surveillance"

**Supplementary Figures**

**Figure S1. Systemic viral infection leads to memory CD8$^+$ T cell populations in SLO and SMG for comparative analysis. A.** Experimental setup to study CD8$^+$ T cell behavior and phenotype in SMG and SLO. GFP$^+$ OT-I CD8$^+$ T cells were i.v. injected on day -1. Recipient mice were infected with LCMV-OVA on day 0 and analyzed for viral titer determination and flow cytometry on indicated days. **B.** Viral titer determination on days 3 and 5 p.i. with LCMV-OVA. Each dot represents one mouse, lines indicate median. **C.** Representative flow cytometry analysis of adoptively transferred GFP$^+$ OT-I T cells after gating on FSC/SSC-H lymphocyte CD45$^+$ singlets. Numbers indicate percentages. **D.** Total and OT-I T cell numbers in spleen PLN and SMG at indicated days p.i. with LCMV-OVA. Each dot represents one mouse, lines indicate median. **E.** Surface marker expression on transferred OT-I T cells at indicated time points. Dotted lines are isotype or FMO (for CD69) controls. **F.** Quantification of cell surface marker expression. Plotted are mean ± SD. Data in B are from 2 independent experiments with a total of 5-10 mice per time point. Data in F are pooled from 3-4 independent experiments with a total of 6 to 15 mice per time point, except CD69 data showing 1 of 2 independent experiments with 3 mice/time point.

**Figure S2. Dynamic motility of P14 CD8$^+$ $T_{PLN-M}$ and $T_{RM}$ before and after cognate peptide injection. A, B.** GFP$^+$ P14 CD8$^+$ $T_{PLN-M}$ and SMG $T_{RM}$ speeds (**A**) and meandering index (**B**) before cognate peptide injection. **C, D**. $T_{RM}$ speeds (**C**) and meandering index (**D**) after injection of irrelevant ($OVA_{257-264}$) or cognate ($gp_{33-41}$) peptide. Red lines indicate median. A and C were analyzed using a Student's t-test and B and D with Mann-Whitney test. Data in A and B are from 4-7 mice and in C and D from 1-4 mice in 1-3 independent experiments.

**Figure S3. Characterization of CD11c-YFP$^+$ macrophages. A.** Gating strategy for macrophage marker staining in SMG of CD11c-YFP mice. **B.** Immunofluorescent SMG section of CD11c-YFP$^+$ signal and Iba-1. Iba-1$^+$ CD11c-YFP$^-$ cells are marked by white arrows. Scale bar, 50 µm. **C.** Immunofluorescent SMG section showing tissue macrophages close to EpCAM$^+$ secretory epithelium (arrow) and SMA$^+$ interstitial vasculature (arrowhead). Scale bar, 40 µm. **D.** Confocal SMG section showing pTyr signal in tissue macrophages. Scale bar, 3 µm. **E.** Quantification of pTyr$^+$ protrusions. Left panel shows macrophage/pTyr/DAPI signal, right panels show three consecutive z-stacks (spacing 1 µm) of macrophage/pTyr signal. **F.** Immunofluorescent LG section in memory phase (≥ 30 days p.i. with LCMV-OVA) showing GFP$^+$ OT-I $T_{RM}$ adjacent to CD11c-YFP$^+$ tissue macrophages (indicated by yellow arrowheads). Scale bars, 1000 (left), 100 (middle) and 20 µm (right). **G.** Examples of colocalization of CD68$^+$ macrophage (red) and CD3$^+$ T cell clusters (brown; encircled) in human parotid salivary gland. Scale bar, 50 µm. **H.** Dispersed T cells (brown) in human parotid salivary gland associate with macrophage cell bodies or thin protrusions (red), indicated by arrowheads. Scale bar, 20 µm.

**Figure S4. Analysis of integrins and $T_{PLN\text{-}M}$ blockade *in vivo*. A.** Expression of integrins on memory phase OT-I T cells in spleen, PLN and SMG. **B.** mAb saturation testing. Immediately after 2PM recording, PLN and SMG single cells suspensions were either incubated with the same blocking mAb, followed by a Cy5-labeled Ab against the first mAb (1$^{st}$ and 2$^{nd}$ Ab), or just with the Cy5-labeled 2$^{ndary}$ Ab (2$^{nd}$ Ab only) and analyzed by flow cytometry. **C.** Frequency distribution of $T_{PLN\text{-}M}$ speeds after integrin blockade. Arrows indicate median (µm/min). **D.** Confocal image of $\beta$1 staining of SMG section. Arrowhead indicates macrophage - $T_{RM}$ interface. Scale bar, 4 µm **E.** Confocal image of E-cadherin staining of SMG section. Scale bar, 5 µm. **F.** Frequency distribution of $T_{PLN\text{-}M}$ speeds after PTx treatment. Arrows indicate median (µm/min). Data in C and F are pooled from 2-5 independent experiments with a total of 2-7 mice and analyzed with unpaired Student's t-test. ***, $p < 0.001$.

**Figure S5. LG $T_{RM}$ motility parameter after macrophage depletion. A.** Frequency distribution of $T_{RM}$ speeds in DTx-treated CD11c-YFP or CD11c-DTR bone marrow chimera (control and macrophage-depleted, respectively). Arrows indicate median (µm/min). **B.** Percent of tracks making U-turn. Bars indicate 95% confidence intervals. Data are pooled from 2-3 independent experiments with 5 mice each and analyzed by unpaired t-test (A).***, $p < 0.001$.

**Figure S6. Tissue macrophage-mediated efferocytosis during viral infection of SMG. A.** Experimental layout for local infection of the SMG via retrograde WD injection. **B.** Immunofluorescent SMG sections at 48 h post WD injection with MCMV-OVAmCherry. Infection is restricted to the SMG. Scale bar, 50 µm. **C.** Quantification of mCherry⁺ foci per FOV in SMG sections 48h after MCMV-OVAmCherry infection in the presence or absence of macrophages and $T_{RM}$. **D.** Intensity of mCherry signal per spot in the presence or absence of $T_{RM}$ in macrophage-depleted SMG. **E**. $T_{RM}$ and tissue macrophages cluster around infected cells. Scale bar, 50 µm. **F**. Macrophage engulfing infected cell (arrow). Scale bar, 20 µm. Data in C were analyzed with Kruskal-Wallis and Dunn's multiple comparisons test and in D with Student's unpaired t-test. ***, $p < 0.001$.

**Figure S7. Proposed model of tissue macrophage-assisted $T_{RM}$ immune surveillance in SMG. A.** Confined SMG $T_{RM}$ display distinct migration modes. *Ex vivo* confined SMG TRM respond to chemoattractants and adhesion molecules, while displaying base-line motility through friction. Furthermore, SMG $T_{RM}$ respond to physical cues from their environment to insert protrusions and move between adjacent structures that are not tightly connected. **B.** By associating with protrusion-forming tissue macrophages, which lack extensive contacts with surrounding tissue cells, $T_{RM}$ exploit their distinct motility modes for organ surveillance. Arrows indicate protrusion direction. E, epithelium; M, tissue macrophage.

**Supplementary movie Legends**

**Movie S1. Intravital imaging of $T_{PLN-M}$ migration in PLN.** $T_{PLN-M}$ (green) display amoeboid shapes while migrating in the T cell zone of PLN close to HEV (red). Scale bar, 20 µm and 10 µm (zoom). Time in min:s.

**Movie S2. Intravital imaging of $T_{RM}$ migration in SMG.** $T_{RM}$ (green) migrate in the SMG between blood vessels (red). $T_{RM}$ frequently display multiple leading edges or shape changes. Scale bar, 20 µm and 10 µm (zoom). Time in min:s.

**Movie S3. Confocal image of SMG $T_{RM}$ colocalization with tissue macrophages.** $T_{RM}$ (green) are in close proximity to tissue macrophages (blue). EpCAM-1 (white) depicts epithelial cells. Close up shows a $T_{RM}$ in a duct, identified by ZO-1 staining (red).

**Movie S4. Intravital imaging of SMG $T_{RM}$ migration along tissue resident macrophages.** Time series (left) and time accumulated overlay (right), including xy and xz projections. $T_{RM}$ (green) frequently move along thin protrusions of $CD11c^+$ tissue resident macrophages (blue). Blood vessels are shown in red. Major ticks 50 µm and 20 µm (zoom). Time in min:s.

**Movie S5. Time-lapse imaging of $T_N$ migration under *in vitro* confinement on CCL21/ICAM-1-coated plates.** $T_N$ (red) show efficient migration on CCL21/ICAM-1-coated glass slides under agarose. Scale bar, 50 µm. Time in min:s.

**Movie S6. Time-lapse imaging of $T_{RM}$ migration under *in vitro* confinement on CXL10 + CXCL12/ICAM-1-coated plates.** $T_{RM}$ (green) show efficient migration on chemokine/ICAM-1-coated glass slides under agarose. Scale bar, 20 µm. Time in min:s.

**Movie S7. Time-lapse imaging of $T_{RM}$ and $T_N$ migration under *in vitro* confinement on HSA-coated plates.** $T_{RM}$ (green) but not $T_N$ (red) show efficient migration on HSA-coated glass slides under agarose. Scale bar, 50 µm. Time in min:s.

**Movie S8. Time-lapse imaging of $T_{RM}$ and $T_{PLN\text{-}M}$ migration under *in vitro* confinement.** $T_{RM}$ (left) but not $T_{PLN\text{-}M}$ (right) show efficient migration on HSA-coated glass slides under agarose. Major ticks 20 µm. Time in min:s.

**Movie S9. High temporal resolution analysis of $T_{RM}$ motility under *in vitro* confinement.** $T_{RM}$ migration on HSA-coated glass slides under agarose. Scale bar, 10 µm. Time in min:s.

**Movie S10. Time-lapse imaging of $T_{RM}$ migration in presence of EDTA.** Left panel. $T_{RM}$ form protrusions but do not translocate under agarose in the presence of EDTA. Right panel. $T_{RM}$ migration is restored within clusters of $T_N$ (red), which serve as scaffold for $T_{RM}$ motility. Major ticks 5 µm. Time in min:s.

**Movie S11. Time-lapse imaging of $T_{RM}$ migration inside $T_N$ cluster in presence of EDTA.** $T_{RM}$ (green) show motility while traversing a cluster of $T_N$ in presence of EDTA. Nuclei are labelled with Hoechst (blue). Major ticks 5 µm. Time in min:s.

**Movie S12. Time-lapse imaging of $T_{RM}$ migration along polystyrene beads in presence of EDTA.** $T_{RM}$ (green) show motility while migrating in contact with polystyrene beads in presence of EDTA. Scale bar, 20 µm. Time in min:s.

**Movie S13. Intravital imaging of $T_{RM}$ migration and association with macrophages after blocking of αL, α4 and αE integrins.** $T_{RM}$ (green) move rapidly and associate with macrophages (blue) 16 h after treatment with integrin-blocking mAbs. Blood vessels are shown in red. Scale bar, 20 µm and 10 µm (zoom). Time in min:s.

**Movie S14. Intravital imaging of $T_{RM}$ migration and association with macrophages after treatment with PTx.** $T_{RM}$ (green) move rapidly and associate with macrophages (blue) 3 h after treatment with PTx. Scale bar, 20 µm and 10 µm (zoom). Time in min:s.

**Movie S15. Time-lapse imaging of $T_{RM}$ and tissue macrophages under *in vitro* confinement.** $T_{RM}$ (green) detaching from SMG macrophages (blue) under agarose. Scale bar, 50 µm. Time in min:s.

**Movie S16. Time-lapse SUSHI of SMG section during hyperosmotic challenge.** Extracellular space (black) at time point “0:00” depicts image before starting at “0:20” with gradual increase of osmolarity to 350 mOsm/L. During challenge, ECS increases, in particular in interstitium. Scale bar, 10 µm. Time in h:min.

**Movie S17. Confocal image of tissue macrophage protrusions and basement membranes in SMG.** $T_{RM}$ (green) are in close proximity to tissue macrophages (blue). EpCAM-1 (white) depicts epithelial cells. Close up shows a tissue macrophage protruding between basement membranes labeled by laminin (magenta).

**Movie S18. Intravital imaging of $T_{RM}$ migration in the absence of macrophages.** $T_{RM}$ (green) show migration in the absence of macrophages but are mostly confined to individual acini and ducts. Stroma is white, and outlines of acini and ducts are indicated in magenta. Time in min:s.

**Movie S19. Intravital imaging of $T_{RM}$ migrate along a macrophage to enter acini.** $T_{RM}$ (green) migrating along a macrophage (blue) to enter an acinus. Stroma is white, and outlines of acini and ducts are indicated in magenta. $T_{RM}$ entering the acinus is indicated with an arrowhead. Second half of the Supplementary Video shows the image sequence in 3D. For better visibility, one acinus has been masked manually. Time in min:s.

**Movie S20. Intravital imaging of $T_{RM}$ migration in the absence of macrophages after 5 days of DTx treatment.** $T_{RM}$ (green) show migration in the absence of macrophages but are confined to individual acini and ducts (white). Time in min:s.

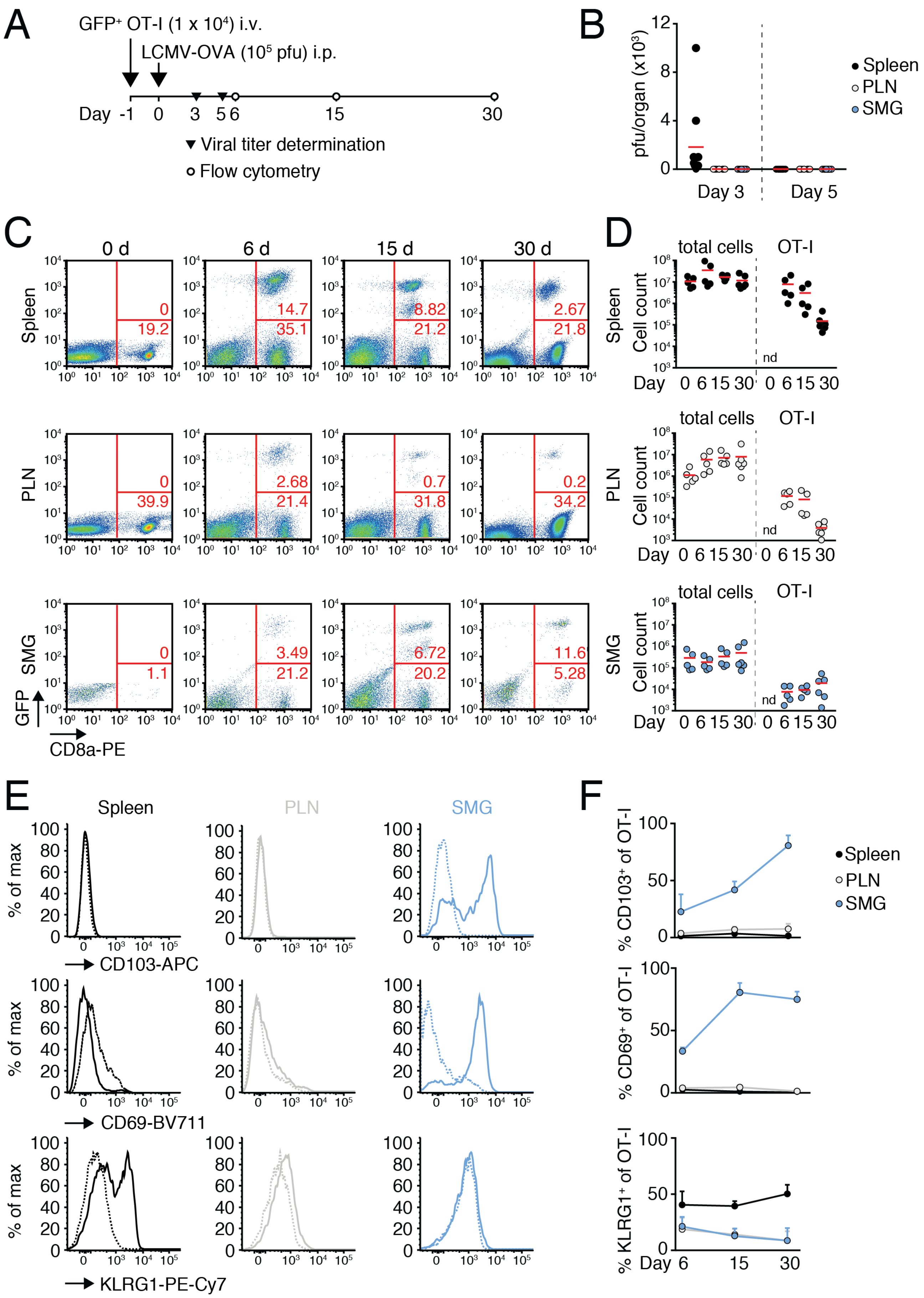

A
GFP+ OT-I (1 x 10^4) i.v.
LCMV-OVA (10^5 pfu) i.p.
Day -1 0 3 5 6 15 30
▼ Viral titer determination
○ Flow cytometry
B
pfu/organ (x10^3)
Day 3
Day 5
Spleen
PLN
SMG
C
0 d
6 d
15 d
30 d
Spleen
PLN
SMG
GFP
CD8a-PE
D
total cells
OT-I
Cell count
nd
Day 0 6 15 30 0 6 15 30
E
Spleen
PLN
SMG
% of max
CD103-APC
CD69-BV711
KLRG1-PE-Cy7
F
% CD103+ of OT-I
% CD69+ of OT-I
% KLRG1+ of OT-I
Day 6 15 30


Figure S1

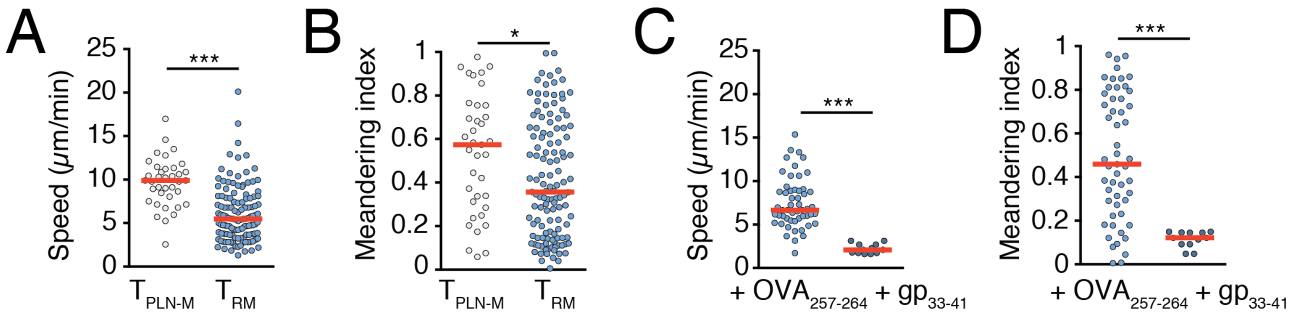

A
Speed (μm/min)
***
T PLN-M
T RM
B
Meandering index
*
T PLN-M
T RM
C
Speed (μm/min)
***
+ OVA 257-264
+ gp 33-41
D
Meandering index
***
+ OVA 257-264
+ gp 33-41


Figure S2

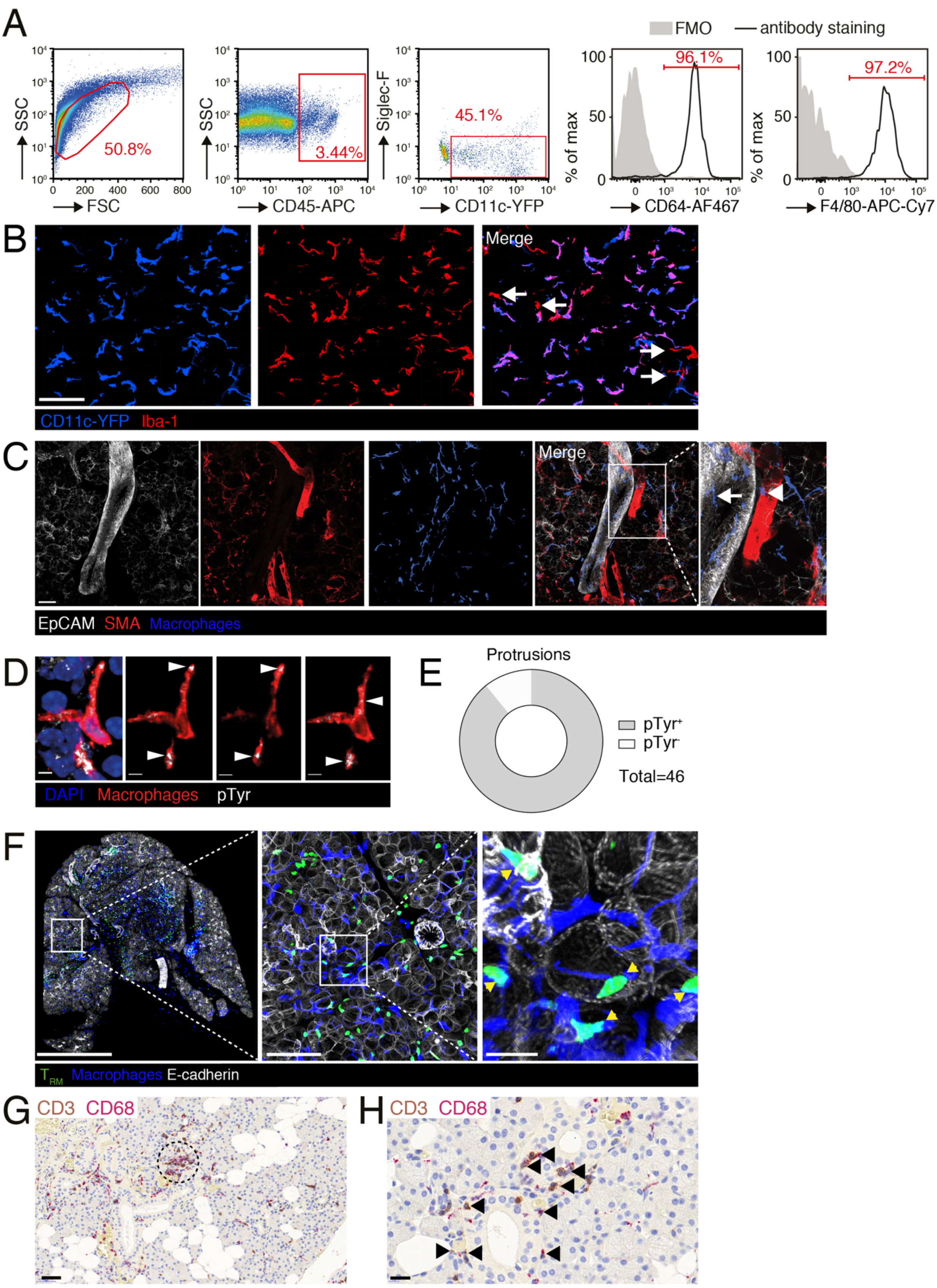


148

Figure S3

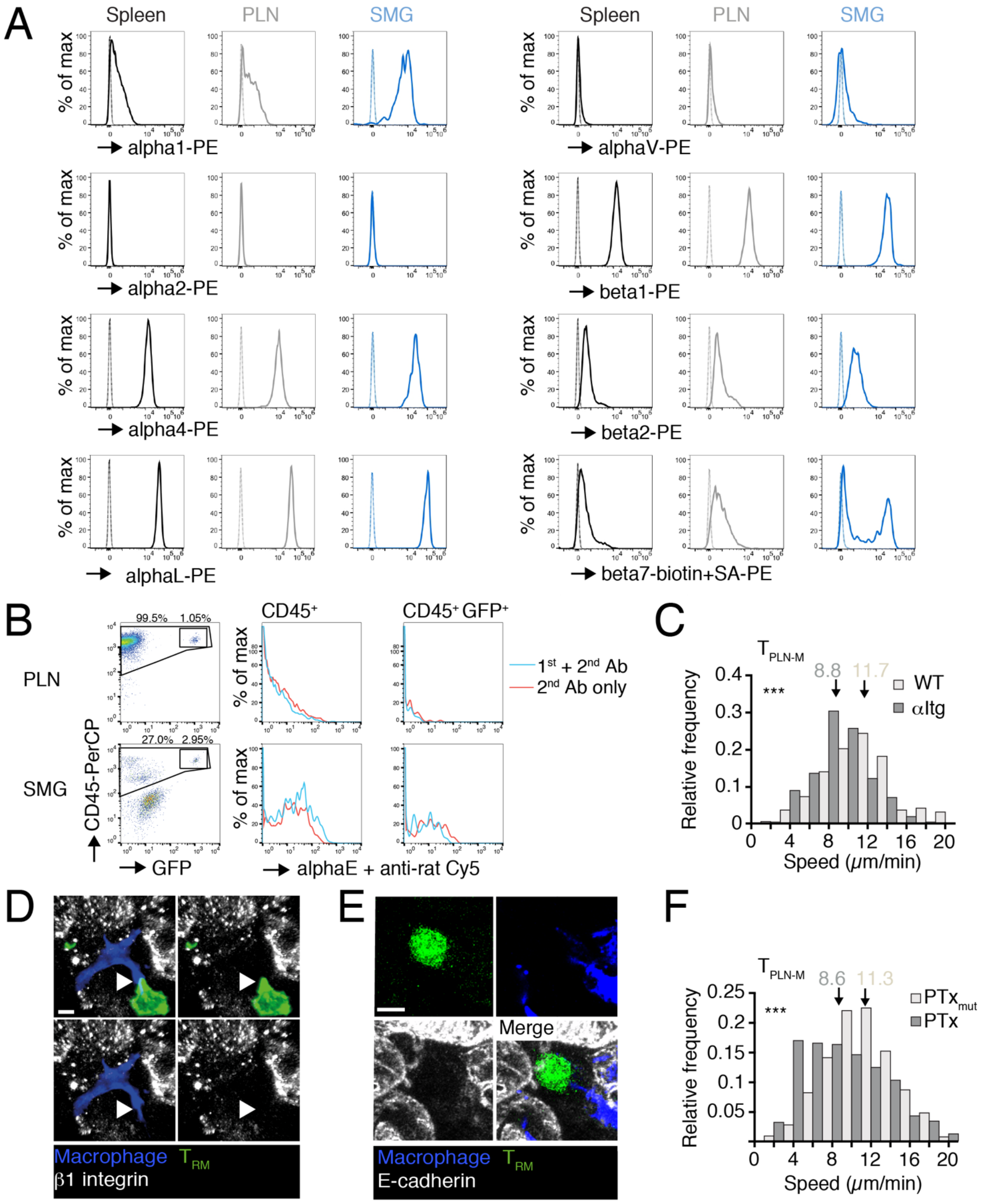


149 Figure S4

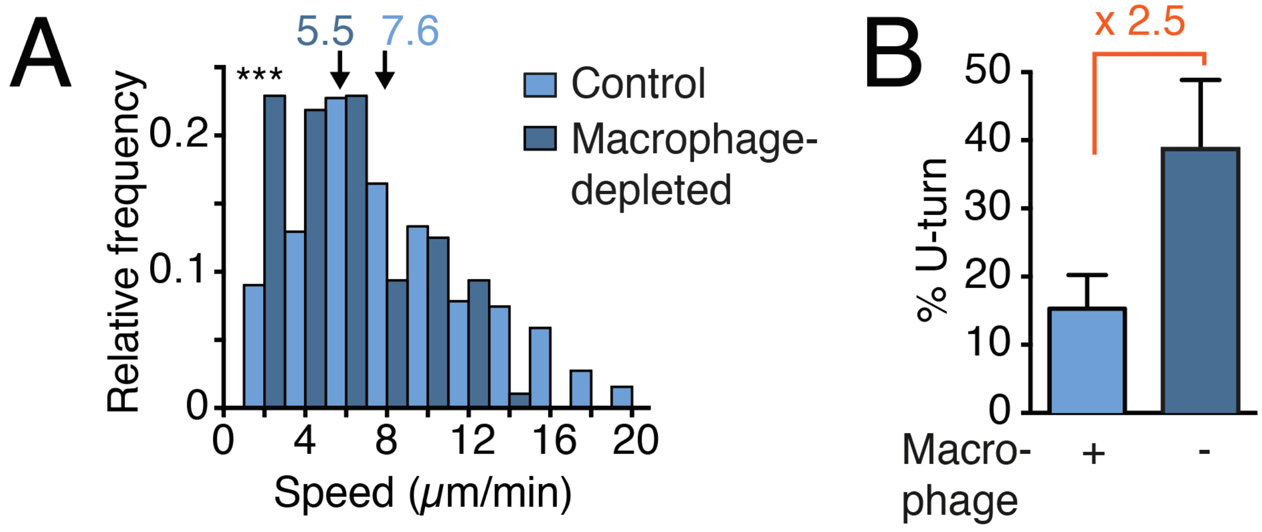


150 Figure S5

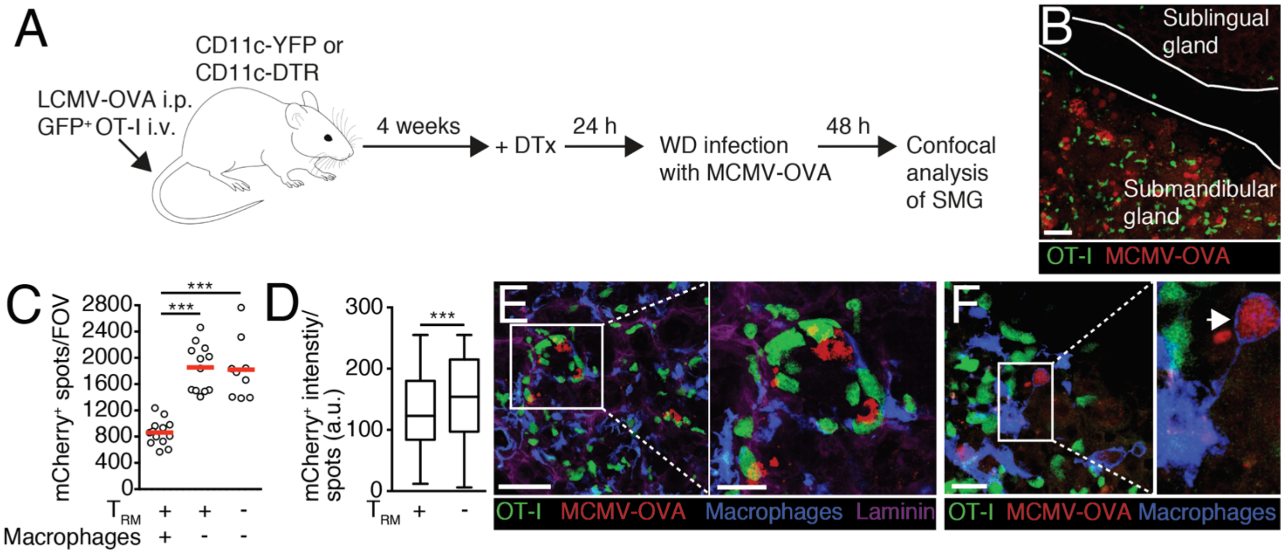


151 Figure S6

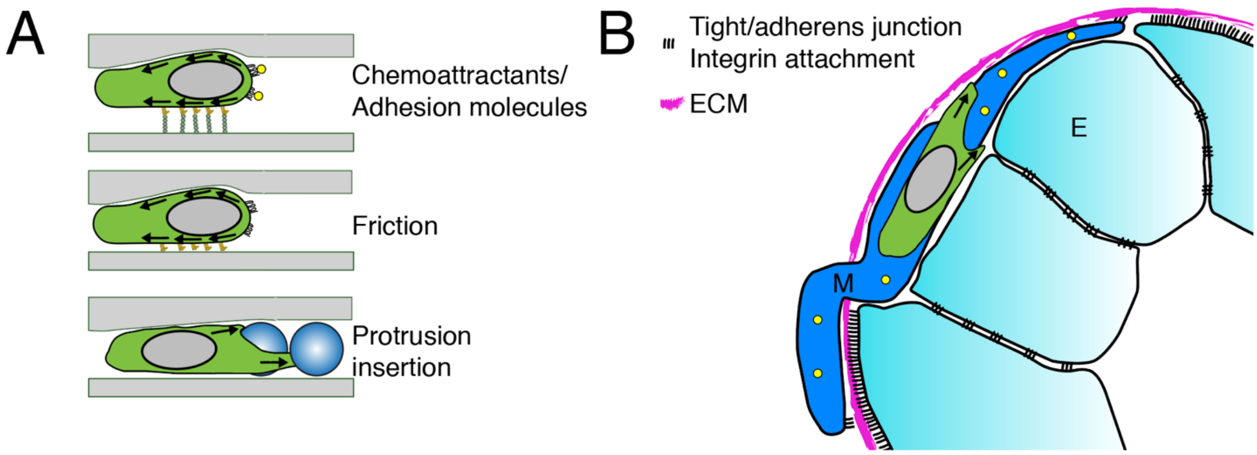


152 Figure S7
